# Anisotropic Articular Cartilage Biofabrication based on Decellularized Extracellular Matrix

**DOI:** 10.1101/2024.07.30.605411

**Authors:** Anna Puiggalí-Jou, Isabel Hui, Lucrezia Baldi, Rea Frischknecht, Maryam Asadikorayem, Jakub Janiak, Parth Chansoria, Maxwell C. McCabe, Martin J. Stoddart, Kirk C. Hansen, Karen L. Christman, Marcy Zenobi-Wong

## Abstract

Tissue-engineered grafts that mimic articular cartilage show promise for treating cartilage injuries. However, engineering cartilage cell-based therapies to match zonal architecture and biochemical composition remains challenging. Decellularized articular cartilage extracellular matrix (dECM) has gained attention for its chondro-inductive properties, yet dECM-based bioinks have limitations in mechanical stability and printability. This study proposes a rapid light-based bioprinting method using a tyrosine-based crosslinking mechanism, which does not require chemical modifications of dECM and thereby preserves its structure and bioactivity. Combining this resin with Filamented Light (FLight) biofabrication enables the creation of cellular, porous, and anisotropic dECM scaffolds composed of aligned microfilaments. Specifically, we investigate the effects of various biopolymer compositions (i.e., hyaluronic acid, collagen I, and dECM) and inner architecture (i.e., bulk light vs FLight) on immune response and cell morphology, and we investigate their influence on nascent ECM production and long-term tissue maturation. Our findings highlight the importance of FLight scaffolds in directing collagen deposition resembling articular cartilage structure and promoting construct maturation, and they emphasize the superiority of biological-rich dECM over single-component materials for engineering articular cartilage, thereby offering new avenues for the development of effective cartilage tissue engineering strategies.

## 1. Introduction

Cartilage tissue has a low self-repair capacity because of its aneural, alymphatic, and avascular characteristics.[1] Despite considerable work, the generation of fibrocartilage and hypertrophic cartilage instead of hyaline cartilage is a common outcome.[2] Cell-based tissue engineering strategies used in the clinical repair of articular cartilage have used hyaluronan (HA) [3], fibrin[4], collagen I (Col I)[5], agarose[6], and polyglactin scaffolds[7], with HA and Col I among the most common. [8] For instance, Biocart^TM^II are scaffolds of freeze-dried fibrin/HA mixtures[3] and examples of Col I include MACI[9], NeoCart^®^[10], Novocart-3D^®^[11], and CaRes®.[8] However, while Col I is present in various tissues, hyaline cartilage contains over 80% type II collagen (Col II).[12] Previous studies have revealed that articular cartilage decellularized extracellular matrix (dECM), mainly composed of Col II, possesses inherent chondro-inductive properties, guiding human mesenchymal stromal cells (hMSCs) to enhance the expression of cartilage-specific genes and secrete glycosaminoglycans (GAGs)-rich matrix. These studies have shown articular cartilage dECM’s potential for regenerating cartilage with reduced hypertrophic characteristics compared to Col I scaffolds.[13,14]

dECM-based materials have been used for articular cartilage repair, as well as to reconstruct mammary tissue[15], skin[16], bone[17], heart[18], kidney[19] and liver[20]tissue. Their highly similar composition to the source tissue can enable better recapitulation of the cell microenvironment, contributing to better tissue maturation.[21] In the past decade, different articular cartilage dECM resins have been investigated to promote hMSCs chondrogenesis.[22–28] Li et al. showed that articular cartilage dECM inhibits hypertrophic differentiation of hMSCs and identified collagen type XI as the ECM component with the most significant effects on enhancing cartilage matrix production and inhibiting its degradation.[14] Lian et al. demonstrated that Col II reduces the expression of collagen type X, which is typically associated with calcified cartilage.[29] It is clear that articular cartilage dECM contains many beneficial components for cartilage regeneration.

To engineer articular cartilage with precise architecture, scientists rely on biofabrication techniques.[30] However, despite significant progress in the tissue engineering field, the existing dECM-based resins face challenges related to their mechanical stability and printability; this limits their use for 3D printing applications. Consequently, various physical and chemical strategies, including nanofiber-reinforced dECM[31,32] and conjugation of methacryloyl functional groups to dECM-based resins (e.g., methacrylated dECM) have been investigated.[33–35] However, functionalizing collagen (the main component of the dECM) may result in the loss of its fibrillar structure, potentially reducing desirable characteristics such as bioactivity and biodegradability. To address these limitations, Kim et al. used ruthenium/sodium persulfate (Ru/SPS) to crosslink cornea-and heart-derived dECM bioresins without chemically modifying the dECM.[36] Given that dECM resins contain proteins rich in tyrosine residues, Ru/SPS system can facilitate rapid and cytocompatible crosslinking of dECM resins. Recently, Lian and colleagues explored dECM resins containing Ru/SPS to volumetrically bioprint heart and meniscal tissue.[37] To our knowledge, however, no photoresins combining articular cartilage dECM with Ru/SPS have been yet developed to repair cartilage defects with mimicking architecture.

Most of the current tissue engineering approaches rely on bulk hydrogels consisting of dense polymeric networks with a nanoscale mesh size, thus constraining encapsulated cells and failing to replicate tissue inner architecture. Recently, the field has been shifting towards hydrogels with higher void, which can increase nutrient diffusion[38], cell proliferation, migration[39,40] and ECM production[41,42], improving overall tissue maturation[38,43–47]. Filamented light (FLight) biofabrication has emerged as a promising technique in this context. [48] It enables the rapid production of macroporous anisotropic constructs within seconds, effectively increasing void space and offering a suitable method for mimicking the inner architecture of anisotropic tissues.[48] Due to a phenomenon known as optical modulation instability (OMI) and the change in refractive index upon crosslinking, FLight generates hydrogels composed of aligned microfilaments and microchannels with diameters ranging from 2 to 30 µm.[48] Such microarchitecture showed excellent cell instructive capabilities in controlling cell and ECM alignment. [48] In contrast to freeze-drying [49], electrospun fibers [50], and other non-cell compatible techniques, FLight technology enables the biofabrication of anisotropic dECM scaffolds in the presence of cells. This alignment is crucial to replicating the architecture of articular cartilage that can mature into constructs with native tissue-like mechanical properties.

hMSCs are used in this study because they are an easily available autologous cell source for regenerative medicine.[51] They are found in various body tissues and play a vital role in tissue repair and regeneration. hMSCs are mesenchymal progenitors capable of differentiating into bone, fat, muscle, and cartilage.[52] These cells are easily cultured and expanded *in vitro*, making them highly valuable for tissue engineering applications. However, a deeper understanding of the specific signals that drive hMSC differentiation is essential, in order to develop regenerative therapies that guide cell fate and tissue development.

To summarize, we developed photoresins containing the most relevant materials in cartilage tissue engineering (HA, dECM, and Col I) and evaluated two main parameters: the effect of their composition and their microarchitecture on hMSC chondrogenic differentiation. We emphasize the significant influence of both biochemical cues and architecture on hMSC chondrogenesis in early stages by nascent metabolic labeling and in late stages by histology (**Figure 1**). Higher void space and the presence of dECM were the most chondroinductive stimuli. Our findings also highlight the preservation and guided alignment of dECM and Col I after FLight printing. Furthermore, we showed the possibility of printing intricate structures within seconds at high resolution, using dECM and Col I-containing resins. Therefore, this study provides crucial insights for optimizing cartilage tissue engineering and advancing regenerative approaches.

**Figure 1.**
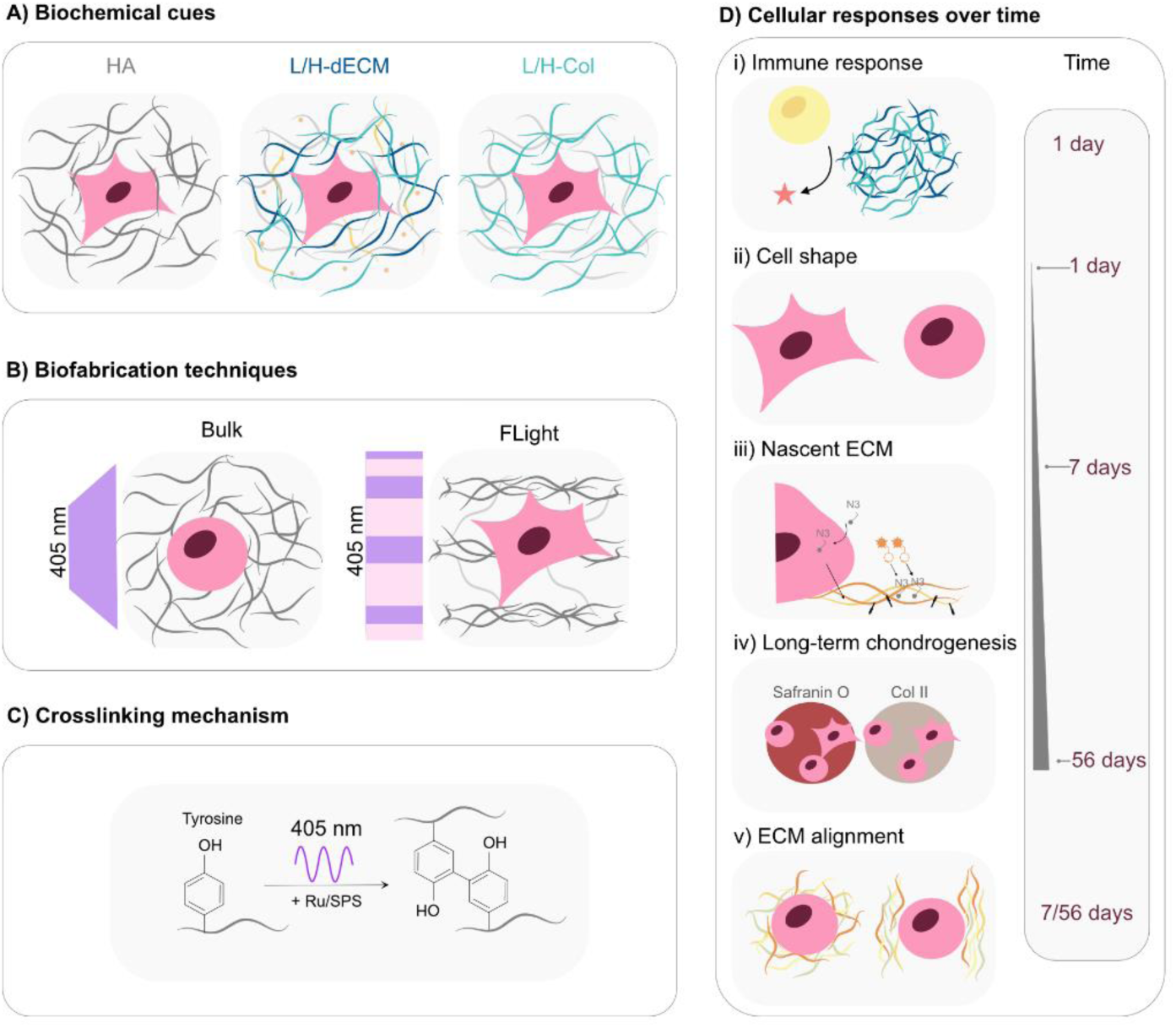
Schematic of the two tested parameters, biochemical cues, and biofabrication methods using Ru/SPS crosslinking mechanism and how they influence the cell response over time. A) Illustration of the different biological cues: HA (HA 0.8%), low (L)-dECM (HA 0.8% + 2.5 mg mL^-1^ dECM), high (H)-dECM (HA 0.8% + 5 mg mL^-1^ dECM), low (L)-Col I (HA 0.8% + 2.5 mg mL^-1^ Col I), and high (H)-Col I (HA 0.8% + 5 mg mL^-1^ Col I). B) Presentation of the biofabrication methods to impart different physical cues: bulk and FLight. C) Schematic of the crosslinking mechanism: upon light irradiation (405 nm), tyrosine groups in L/H-dECM, L/H-Col I, and tyrosine-modified HA are oxidized by Ru/SPS, forming tyrosyl free radicals that subsequently create covalent di-tyrosine bonds with nearby tyrosine moieties. D) Representation of different cellular responses: immune response with THP-1 cells, hMSC shape after 1 day of incubation in chondrogenic media, nascent ECM production in chondrogenic media after 1 and 7 days of culture, long-term chondrogenesis (after 56 days), and quantification of aligned ECM produced by hMSCs at different time points (7 and 56 days).

## 2. Results

### 2.1 Preparation and characterization of cartilage photoresins

To prepare biologically relevant resins for articular cartilage repair [44,53] and study their effect on hMSC chondrogenic differentiation; we designed the following five combinations containing HA, dECM at low (L) and high (H) concentrations, and Col I at L and H concentrations: HA (HA 0.8%), low (L)-dECM (HA 0.8% + 2.5 mg mL^-1^dECM), high (H)-dECM (HA 0.8% + 5 mg mL^-1^dECM), low (L)-Col I (HA 0.8% + 2.5 mg mL^-1^Col I), and high(H)-Col I (HA 0.8% + 5 mg mL^-1^ Col I). HA was functionalized with tyramine groups for crosslinking and was present in all combinations to provide structural integrity and avoid shrinkage during culture. HA tyramine was synthesized according to the protocol of Darr et al. [54], achieving a degree of substitution (DS) of 8.8% (**Figure S1**). To isolate ECM from articular cartilage, cartilage was dissected from bovine knees, subjected to acid decellularization, and subsequently digested with pepsin (**Figure 2Ai**). Various decellularization protocols are available [55]; however, we followed a hydrochloric acid–based strategy as outlined by Schneider and Nurnberger [56]. This approach was chosen for its efficacy in DNA removal and avoidance of detergents, which can be challenging to eliminate completely. Following decellularization, the DNA content was measured and found to be below 50 ng DNA mg^-1^ dry tissue (**Figure 2Aii**), thereby meeting the literature-approved limit[57] while preserving half of the original concentration of µg GAGs mg^-1^ dry tissue (**Figure 2Aiii**). Proteomics was used to further characterize the protein composition of the dECM (**Figure 1Aiv-vi**). As expected, collagens were the main component (55 ± 5% total identified intensity); amongst fibrillar collagens, COL2A1 was the most abundant (**Figure 2Av**), and COL9A2 was the most abundant non-fibrillar collagen (**Figure 2Avi**). Interestingly, among the secreted factors (0.53 ± 0.09% total intensity), TGFβ1 was detected, which is a growth factor shown to enhance chondrogenesis (**Figure S2**).

**Figure 2.**
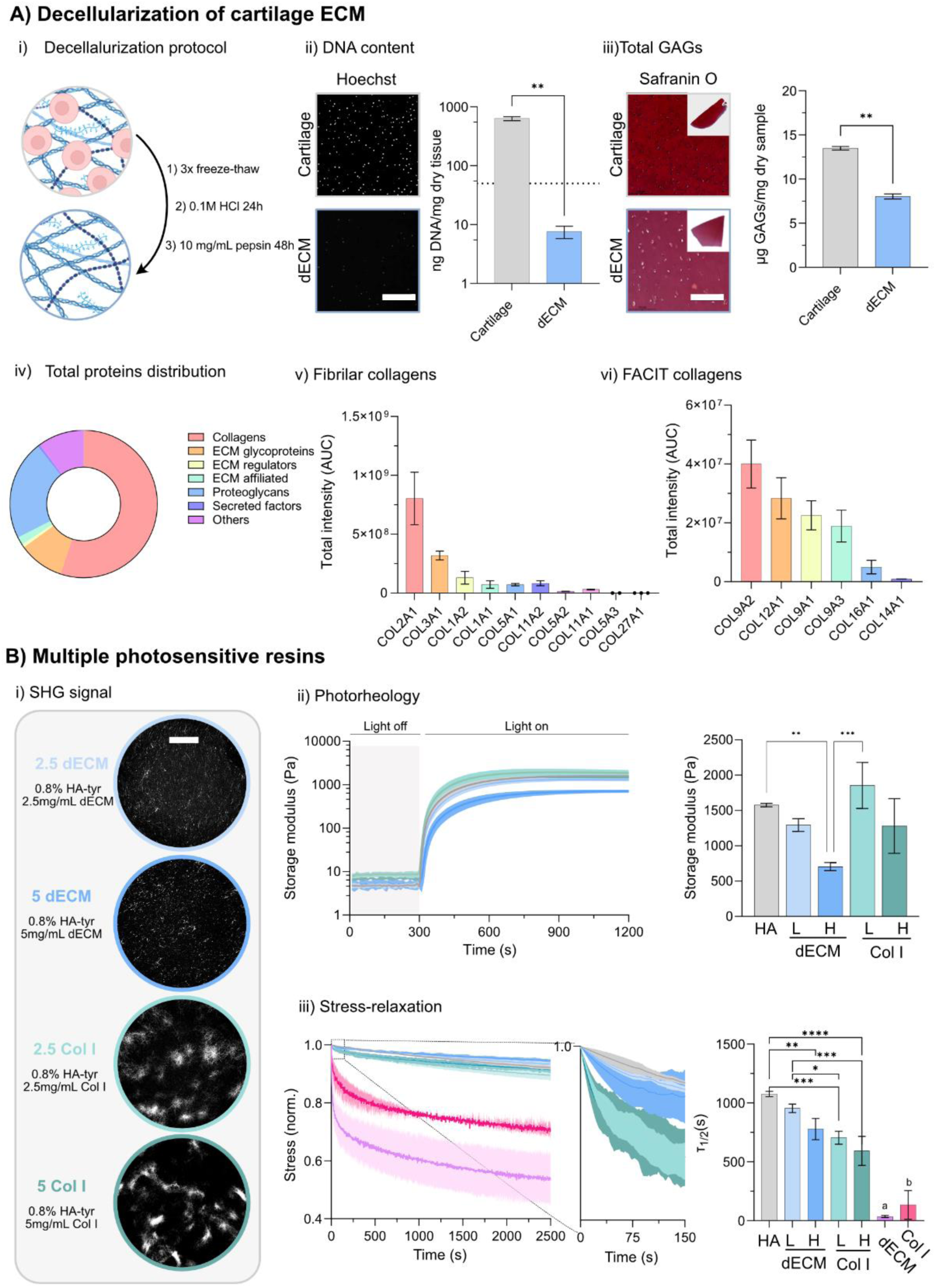
Preparation of biomaterials for light printing. A) Decellularization of articular cartilage ECM. i) Schematic of the decellularization protocol. Figures partially created with BioRender.com. ii) Measurement of DNA content: Hoechst staining before and after the decellularization procedure and graph showing DNA quantification performed with Picogreen Assay. Scale bar = 100 µm. iii) Total GAGs visualized with Safranin O staining before and after the decellularization procedure (scale bar = 100 µm), along with quantification using Blyscan Assay. iv) Global ECM proteomics on three different bovine articular cartilage donors quantifying ECM and ECM-related proteins relative to total ECM area-under-the-curve (AUC) intensity. Total AUC intensity of fibrillar (v) and FACIT (fibril-associated collagens with interrupted triple-helices) collagens (vi). B) Photosensitive resin design. i) Visualization of collagen fibers with second harmonic generation (SHG) of the different gel compositions (L-dECM, H-dECM, L-Col I, H-Col I). Scale bar = 100 µm. ii) Photorheology of the different inks with 0.5 mM Ru/0.05 mM SPS. Light exposure (405 nm) was turned on after 5 min and turned off after 15 min. On the right is their corresponding storage modulus. iii) Stress relaxation curves of hydrogels at a strain of 10% normalized to the initial stress and the corresponding relaxation half-time. Data are shown as means ± SD; three independent experiments; n = 3. Statistical significance was determined using t-test for 1A ii) and iii) and One-way ANOVA with a Tukey’s multiple comparisons test for 1B ii) and iii) (∗p < .05, ∗∗p < .01, ∗∗∗p < .001, and ∗∗∗∗p < .0001, a and b p< .0001 against all other conditions).

As previously demonstrated, the initial stiffness and pore size of hydrogels play a crucial role in governing the maturation of engineered cartilage tissue. [58] Specifically, an initial storage modulus of 1-2 kPa was identified as optimal, leading to a significant stiffness increase over the culture period and ultimately reaching the range of native cartilage (reported in the literature to span from 0.1 to 6.2 MPa).[58] Consequently, we adjusted the different resin formulations (HA, L-dECM, H-dECM, L-Col I, and H-Col I) using photorheology to achieve an initial stiffness of ∼1 kPa (**Figure 2Bii**). To investigate the impact of the biofabrication technique on fibrillogenesis, second harmonic generation (SHG) images were captured after a 30-minute incubation at 37°C for L-dECM, H-dECM, L-Col I, and H-Col I (**Figure 2Bi**); HA did not give any positive signal. Collagen molecules self-assemble into fibrillar structures at a physiological temperature, and electrostatic and hydrophobic bonds hold the fibrillar network together.[59] As expected, image analysis showed a significant difference in the amount of collagen fibers and fibrils among different hydrogels; the amount was higher in L-Col I and H-Col I conditions.

Another important finding is that the addition of ECM components in groups L-dECM, H-dECM, L-Col I, and H-Col I resulted in higher viscoelastic behavior when compared to HA, due to the presence of collagen fibrils. Collagen fibrils possess weak interactions (electrostatic and hydrophobic bonds) that become unbound under stress, allowing for fiber slippage. [60,61] For H-dECM, L-Col I, and H-Col I, the stress relaxation time (τ1/2) of hydrogels decreased significantly (**Figure 2B iii**), and the effect was most pronounced for H-Col I. dECM and Col I alone exhibited even lower τ1/2. Recent studies have found that viscoelasticity in synthetic hydrogels used as cell culture substrates can influence cell behavior, including spreading, proliferation, and differentiation.[33,62,63] Taken together, these mixtures show different viscoelastic behaviors independent of the initial storage modulus. By employing the Ru/SPS crosslinking mechanism, we circumvented chemical modifications of dECM and Col I, thereby preserving their fibrillar structure during photocrosslinking.

### 2.2 Characterization of hydrogels fabricated with bulk photocrosslinking vs FLight

In addition to investigating various resins that introduce distinct biological cues to the system, we conducted a comparative analysis of two fabrication mechanisms designed to impart physical cues to the hydrogels: bulk crosslinking (achieved with LED light) and FLight (**Figure 3A**). FLight fabrication is a light-based technique that rapidly generates anisotropic constructs with embedded microfilaments. A key new feature of the custom-made FLight device is the ability to print directly into 24-well plates, which reduced both the need to manipulate the constructs and the printing time. Previous FLight applications have utilized photosensitive resins based on both chain-growth (e.g., gelatin methacryloyl, hyaluronic acid methacrylate, and alginate methacrylate) and step-growth polymerizations (e.g., gelatin norbonene, and hyaluronic acid norbonene).[45,48] While all these resins have demonstrated the ability to produce anisotropic constructs, they did not fully mimic the chemical complexity of the native tissue environment. We introduced dECM and Col I here, which are more biologically relevant, but their use can be challenging due to their thermal gelation, opacity, and poor shape stability. By using Ru/SPS to induce photocrosslinking, we maintained the collagen structure without reducing cell viability (**Figure S3**). It was notable that after 24 hours in solution, the photoinitiators were washed out of the hydrogels (**Figure S4**).

**Figure 3.**
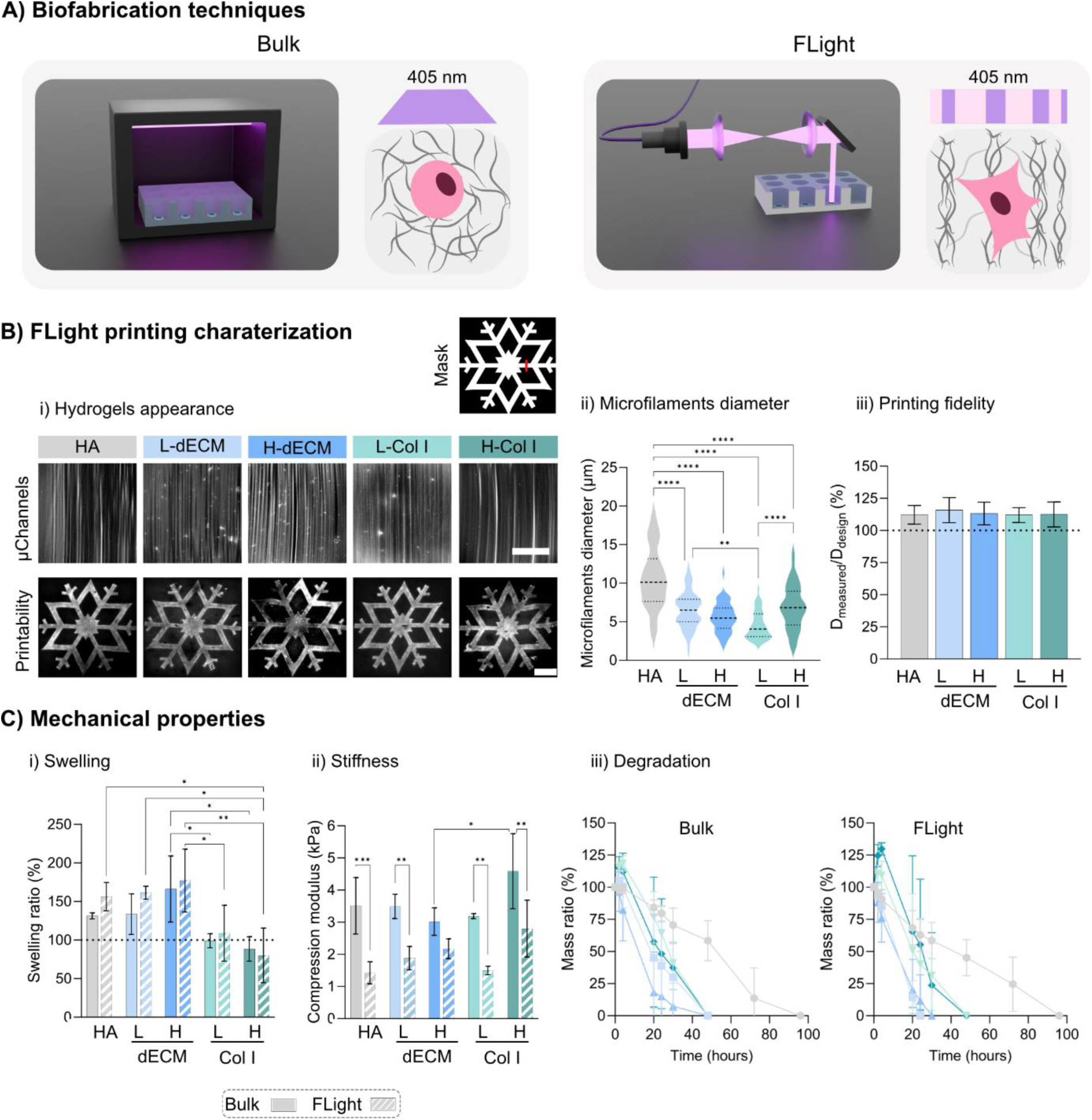
Bulk vs FLight hydrogel fabrication and characterization. A) Illustration of light-based technologies for crosslinking the hydrogels (bulk and FLight). B) FLight printing characterization. i) Microarchitecture characterization of FLight hydrogels with different resins (Scale bar = 50 µm), accompanied by a comparison of the printability of a complex shape: snowflake (top right: pattern design with a red line showing where the printing fidelity was measured) (Scale bar = 1 mm). Measurement of microfilament diameters (ii) and printing fidelity (the black and white projected image is on the top right, and the red line indicates where the measurements were taken to evaluate the printing fidelity) (iii). C) Mechanical properties of bulk (solid) vs FLight (Striped) printed hydrogels: i) Swelling ratio, ii) compression modulus, and degradation profile (iii) of bulk (left) and FLight hydrogels (right). Data are shown as means ± SD; three independent experiments; n = 3. Statistical significance was determined using One-way ANOVA with a Tukey’s multiple comparisons test for 2B ii) and iii) and Two-way ANOVA with a Tukey’s multiple comparisons test for 2C i) and ii) (∗p < .05, ∗∗p < .01, ∗∗∗p < .001, and ∗∗∗∗p < .0001).

To examine the microarchitecture of the FLight hydrogels vs bulk hydrogels, we incorporated rhodamine-acrylate into the resins before printing. For bulk samples, where LEDs serve as non-coherent light sources without exhibiting speckle patterns, no microfeatures resulting from self-focusing phenomena were detected. Only a uniform rhodamine signal was observed, suggesting the presence of a nanoporous polymeric network (**Figure S4**). As expected, highly aligned microfilaments were formed in all FLight constructs (**Figure 3Bi, top**) in the range of 5-10 μm diameter (**Figure 3Bii**), revealing the existence of microchannels capable of enhancing nutrient diffusion and guiding the deposition of cell-secreted ECM.

We further fabricated large, geometrically complex structures using FLight technology (**Figure 3Bi, bottom**) to assess the printing fidelity with the different resins. We printed a snowflake structure with details down to 150 μm and compared it with the initial design (**Figure 3Bi**). All photoresins were able to reproduce the designed structure, but swelling was observed in all groups, as the measured diameter was always larger than the intended design. When we compared the diameter of the thinner strut to the diameter of the printed design, we observed that the strut diameter in L-dECM and H-dECM was slightly higher than in the original design due to swelling (**Figure 3B iii**). H-dECM and H-Col showed more inhomogeneities during printing, likely attributed to the presence of small particles and alterations in light scattering, even though the initial resin transparency was very high (**Figure S4**). Furthermore, removing all uncrosslinked resin was more difficult for H-dECM and H-Col I. Recent studies have demonstrated that this issue can be overcome by incorporating iodixanol into the photoresins.[64,65] However, adding iodixanol can reduce the gels’ compressive modulus, thereby altering their physical properties.[66]

FLight and bulk hydrogels were successfully fabricated using the different photoresins, and the swelling ratios, compressive modulus, and degradation of the crosslinked hydrogels were evaluated as indicators of hydrophilicity and crosslinking density. L-dECM and H-dECM exhibited a significant increase in mass swelling ratios, contrasting with a slight reduction observed for L-Col I and H-Col I in both FLight and Bulk hydrogels (**Figure 3Ci**). This suggested the presence of a stiffer polymer network in hydrogels containing Col I, a finding supported by compression modulus measurements (**Figure 3Cii**). Interestingly, significant differences were observed between FLight and bulk in compression modulus, which were consistently lower for FLight hydrogels due to the higher void space (∼40%). The degradation profile in a physiologically relevant hyaluronidase and collagenase enzyme solution (**Figure 3Ciii**) was influenced by the gel composition but not by the architecture, with L-dECM and H-dECM degrading faster than L-Col I, H-Col I and HA hydrogels. The higher degree of swelling facilitates faster enzymatic cleavage due to better accessibility of the target groups. Fast degradation would ensure fast remodeling of the hydrogel by the encapsulated cells.

### 2.3 Matrix composition and architecture influence immune response

The rationale for using dECM in tissue engineering lies in the advantages of employing a natural biomaterial that mimics the complex tissue-specific microenvironment. The decellularization process removes immune components such as cells and DNA while preserving the protein composition, thereby facilitating cell growth and proliferation.[67] However, residual immunogenic components within dECM bioscaffolds may potentially cause cytocompatibility issues and trigger proinflammatory responses. Investigating the immunogenicity of hydrogels is relevant for future biological applications.

We assessed hydrogel immunogenicity using THP-1-derived macrophages[68], which revealed differential responses based on hydrogel composition and architecture. The THP1-Dual cells feature two inducible reporter genes: Secreted embryonic alkaline phosphatase (SEAP) and Lucia luciferase. SEAP monitors nuclear factor kappa B (NF-κB) pathway activity in the media, while Lucia luciferase tracks the interferon regulatory factors (IRF) pathway. The THP1-Dual cells were seeded on the different hydrogels for 24 hours, either without stimulation or in the presence of 100 ng/mL^-1^ lipopolysaccharide (LPS). LPS is used as a control to elicit an immune response, serving as a benchmark for evaluating the immunomodulatory effects. **Figure 4** demonstrates that L-dECM and H-dECM in bulk hydrogels without stimulation activated the IRF (**Figure 4A**) and NF-κB (**Figure 4B**) pathways. However, when using aligned hydrogels (FLight), there was a significant response reduction. Similar behavior was observed with HA hydrogels for the NF-κB pathway. Interestingly, smaller differences were observed in L-Col I and H-Col I hydrogels, likely due to thermal crosslinking (fibrillogenesis), which can hinder the presence of microchannels. In the presence of LPS, both IRF and NF-κB pathways were activated, as expected, and similarly for all hydrogels.

**Figure 4.**
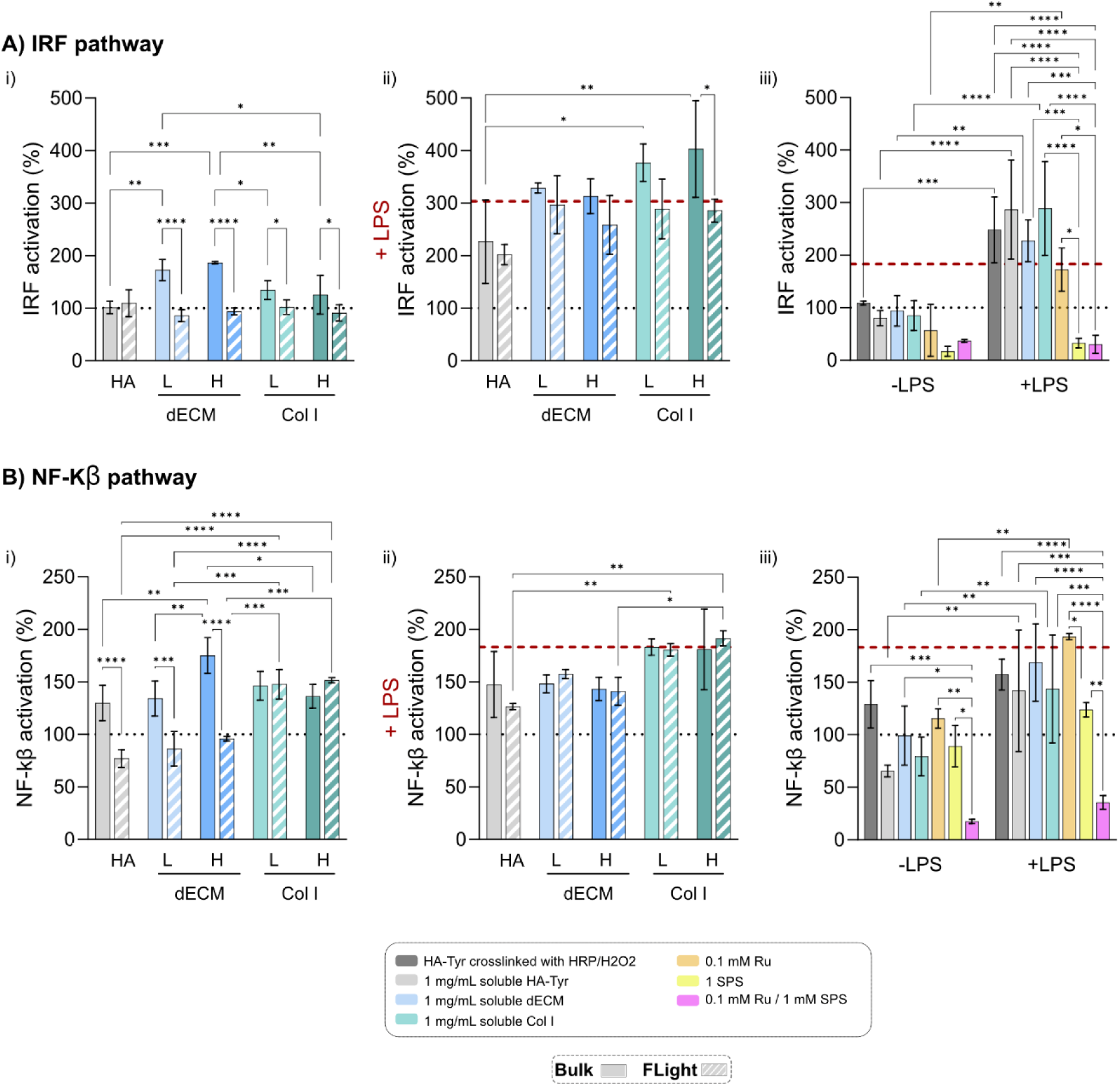
*In vitro* study of macrophages in contact with hydrogels. A) Luciferase readout for IRF activation, B) SEAP reporter readout for NF-κB in THP-1 cells. A and B are the results of cells seeded on hydrogels (i), seeded on hydrogels with the addition of LPS (ii), or including soluble polymers in the media without and with LPS (iii) after 24 hrs. The black dotted line represents the signal from THP-1 cells seeded on the well plate without any stimulant, while the red line represents the signal when LPS is added to the medium. Data are represented as mean ± standard deviation. Statistical significance was determined using two-way ANOVA with a Tukey’s multiple comparisons test (∗p < .05, ∗∗p < .01, ∗∗∗p < .001, and ∗∗∗∗p < .0001) (n = 3 replicates).

Furthermore, to investigate the influence of the individual materials and the crosslinking method on the induction of an immune reaction, HA crosslinked with HRP/H2O2, soluble biopolymers, and Ru/SPS dissolved directly in the media were evaluated. No significant differences were observed between HA, dECM, and Col I soluble in solution. Moreover, using HRP/H2O2 as a crosslinking agent did not alter the results obtained when using Ru/SPS, suggesting that the crosslinking chemistry did not impact the differentiated THP-1 cells. However, reduced activity was observed for pure SPS in the media, probably because SPS has a cytotoxic effect on immune cells. Indeed, Major and colleagues [69] showed that with 1 mM SPS, cell viability was drastically reduced to 20%. Nevertheless, as shown in **Figure S3 and Figure S4**, the designed hydrogels did not retain Ru/SPS after washing, and the viability of the hMSCs was higher than 80%.

These results suggest that the decellularization protocol used in this work appears to be successful in terms of lowering the immune reaction by removing the DNA. Residual DNA can activate the host immune response to dECM. [67] If double-stranded DNA accumulates in the cytosol, type-I interferon (IFN-I) is induced. [70] The IRF pathway was not activated without the addition of LPS, except in the case of bulk dECM hydrogels. However, as the dECM in solution and dECM FLight hydrogels did not elicit an immune response, the reason for the activation probably does not lie in the presence of residual DNA but rather in the internal structure of the hydrogels. Indeed, the hydrogels’ porosity, size, shape, and physicochemical properties can impact the immune response.[71] Macrophages adapt to the characteristics of their surrounding microenvironment, and accumulating evidence points to the crucial influence of physical signals in regulating macrophage activation. [72] Liu and colleagues [72] showed that the spatial confinement of macrophages in scaffolds with cell-sized pores reduces the inflammatory response. Schoenenberger and colleagues [73] noticed that M0 macrophages exposed to aligned polycaprolactone nanofiber substrates for 24 h revealed more elongated shapes associated with pro-reparative (M2) phenotype. Overall, these preliminary results show that FLight hydrogels are non-immunogenic and have potential for *in vivo* applications.

### 2.4 Matrix composition and architecture regulate cell morphology and actin organization

Regulating cytoskeletal architecture is crucial for directing hMSCs towards adipocyte, osteoblast, or myogenic lineages. [74,75] Disruption of actomyosin contraction alone can hinder mesenchymal condensation, thus blocking chondrogenic differentiation.[76] This modulation of hMSC fate and morphogenesis is intricately controlled by dynamic cytoskeletal changes influenced by the microenvironment.

It is therefore interesting to investigate the effects of matrix composition and hydrogel architecture on hMSC morphology. We encapsulated hMSCs in the previously described 3D systems (HA, L-dECM. H-dECM, L-Col I and H-Col I) with bulk and FLight fabrication technique. Adhesion-mediated signaling at the leading edge drives changes in cell morphology and filopodia formation, which enable interactions with the ECM environment. Single-cell images of hMSCs after one day in 3D hydrogels revealed cells with varying morphologies (**Figure 5A**). Quantification of filopodia showed that cells in hydrogels containing L/H-dECM exhibited a higher number of filopodia per cell (**Figure 5Bi**), which were also longer (**Figure 5Bii**). Significant differences were observed between FLight and bulk gels when using HA or L-Col I, where the presence of µ-channels facilitated filopodia protrusions. These results highlighted the importance of hydrogel physical and biological cues on cell morphology and spreading via filopodia.

**Figure 5.**
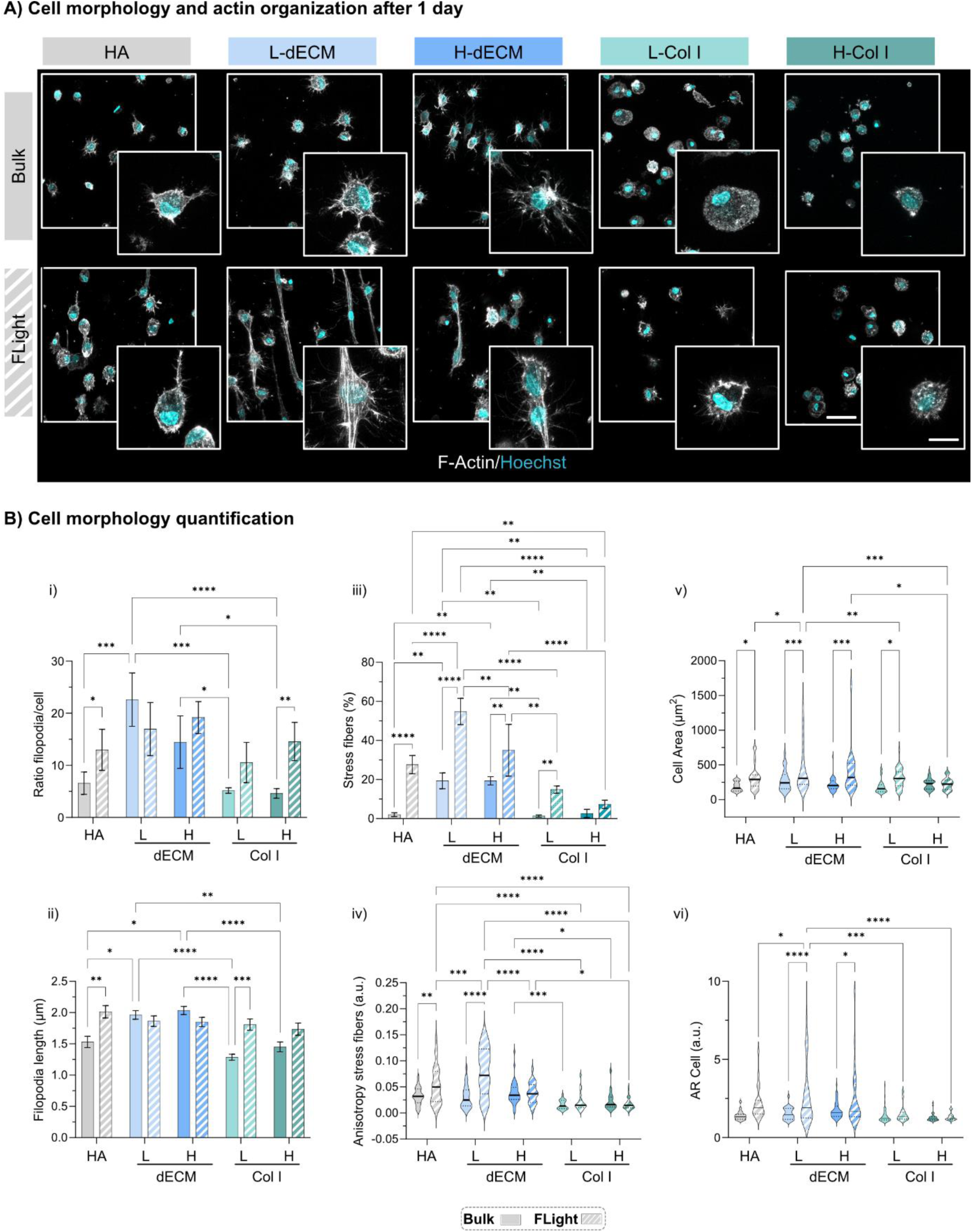
Morphology of hMSCs when encapsulated in 3D hydrogels. A) Representative images of F-actin staining after 24 hours. Scale bars: 20 μm and inset 10 μm. B) Cell morphology analysis, including quantification of filopodia number per cell (i) and maximum filopodia length (ii), percentage of cells forming stress fibers (iii), actin organization and polarization via anisotropy ratios (iv), cell area quantification (v), and aspect ratio (AR) (vi). Data are represented as mean ± standard deviation. Statistical significance was determined using two-way ANOVA with a Tukey’s multiple comparisons test (∗p < .05, ∗∗p < .01, ∗∗∗p < .001, and ∗∗∗∗p < .0001) (n = 3 replicates).

We then investigated whether matrix composition and organization regulated actin organization in hMSCs. Particularly, we examined how the cytoskeleton adapts to the matrix and architecture after one day of chondrogenic culture. Confocal characterization showed more prominent actin stress fibers in dECM gels, while actin staining appeared more granule-like in the other compositions. Notably, there was a significantly higher percentage of cells exhibiting stress fibers in FLight gels than in bulk hydrogels in all biomaterials (**Figure 5Biii**). In addition, quantifying the actin anisotropy ratio demonstrated that stress fibers in FLight hydrogels were more aligned, suggesting the presence of more fibrillar actin (**Figure 5Biv**). Similar results were found when evaluating overall cell area and aspect ratio (AR). The cell area and AR were larger in FLight hydrogels made of L/H-dECM (**Figure 5Bv**) (**Figure 5Bvi**).

Generally, a spreading MSC morphology inhibits chondrogenesis [77,78], while a spherical cell shape initiates chondrogenic differentiation, facilitated by a depolymerized actin cytoskeleton and upregulation of chondrogenic regulators like Sox-9 [79,80]. Therefore, a round morphology resembling chondrocyte morphology might be beneficial. However, recent studies in 3D environments show conflicting results, with some suggesting that round morphology only aids in the initiation, not long-term commitment, of chondrogenesis in MSCs. [81,82] Huang et al. showed that as chondrogenic differentiation proceeds, the materials facilitating cell spreading, which are more viscoelastic, were more conducive to anabolic gene expression and cartilage matrix deposition in the long-term.[82] Lee et al. found that faster-relaxing alginate hydrogels promoted chondrocyte matrix deposition, while slower-relaxing ones limited cell volume expansion, leading to increased interleukin-1β secretion, causing cartilage matrix degradation and cell death. [83] Taken together, these results show that L/H-dECM and FLight architecture promote actin polymerization and alignment. These systems appeared to be more permissive to changes in cell morphology and area, which might be beneficial for cell anabolic activity and long-term commitment to chondrogenesis in the case of hMSCs. [82]

### 2.5 Matrix composition and architecture regulate nascent proteins and proteoglycans

The ECM of mature articular cartilage is rich in proteins and proteoglycans, primarily collagen and aggrecan. [84] After observing short-term differences in cell morphology, we sought to examine whether these differences are translated to ECM production. It is essential to note that cells may only be in contact with the scaffold for a few hours; afterwards, they interact directly with their own newly synthesized matrix. Metabolic labeling enables the visualization of a high-resolution cell-to-cell heterogeneity of the secreted matrix, with information also on its spatiotemporal nature. The method used in this study, pioneered by Loebel and colleagues, involved introducing azide-modified amino acids or sugar analogs into the cell culture media to facilitate their incorporation into nascent matrix components during synthesis. [84,85]

To this end, hMSCs were encapsulated within the previously described five different resin compositions and crosslinked in bulk or FLight matrices. The constructs were cultured in chondrogenic media with continuous presence of either azidohomoalanine (AHA) or N-azidoacetylmannosamine (ManNAz) for 1 or 7 days to visualize nascent proteins (i.e., collagen type II and collagen type VI) or proteoglycans (i.e., aggrecan core protein and its associated chondroitin sulfate residues) (**Figure 6A**). After only 1 day of culture, hMSCs were already surrounded by a nascent protein and proteoglycan layer. After extending the culture period to 7 days, an increase in the thickness of the protein and proteoglycan layer was observed, indicating accumulation of the nascent matrix in the pericellular space (**Figure 6B**). In all conditions, accumulated matrix thickness after 1 day was higher for nascent proteoglycans (0.9 ± 0.2 μm) compared to proteins (0.5 ± 0.2 μm), consistent with observations by Loebel and colleagues. However, after 7 days, deposited proteins (2.6 ± 0.6 μm) were more abundant than proteoglycans (1.8 ± 0.4 μm) (**Figure S5**).

**Figure 6.**
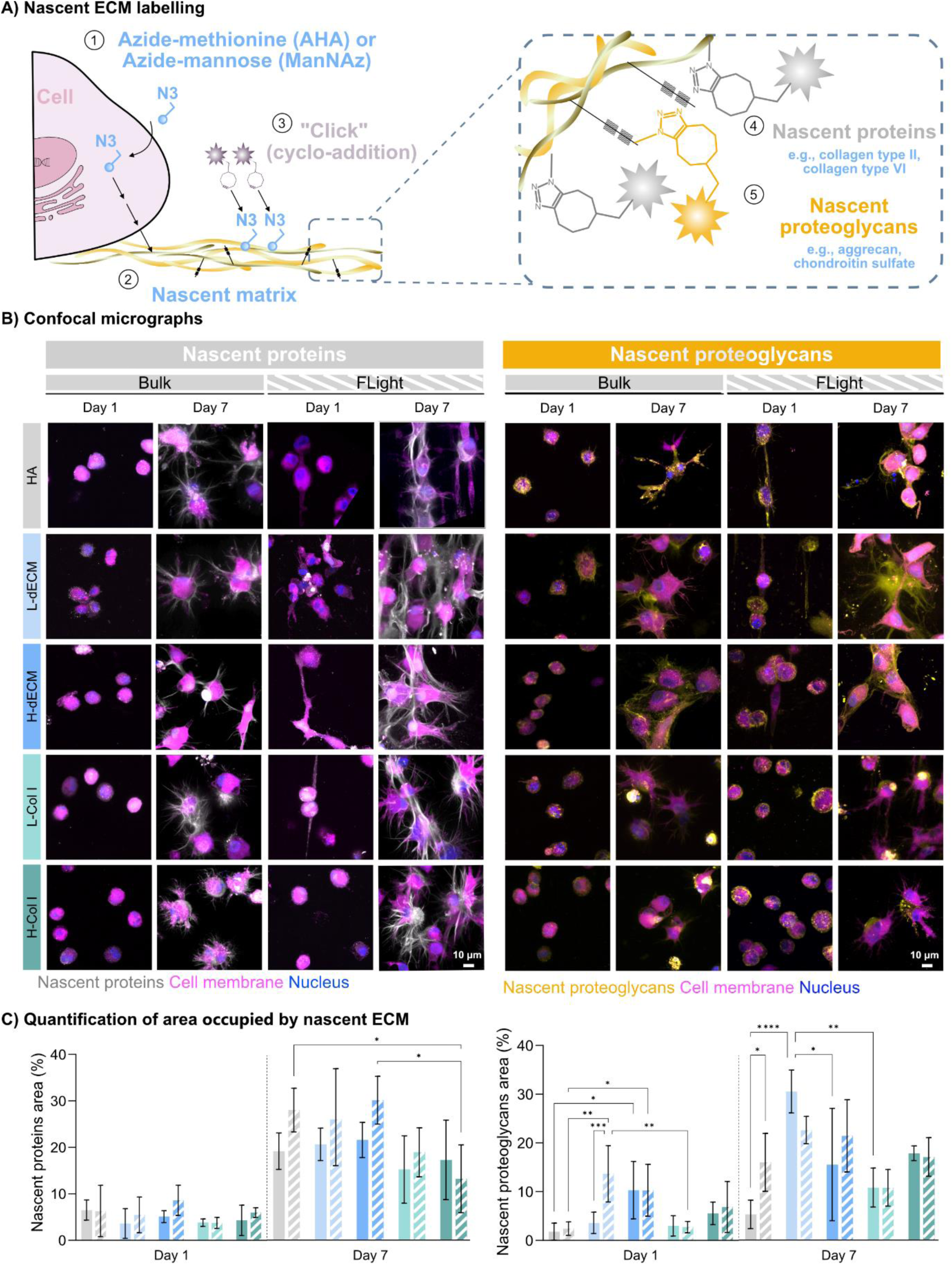
Nascent ECM labelling. A) Schematic of labeling: 1) FLight and bulk gels with hMSCs (5 M cells mL^-1^) were cultured for 7 days with azide-modified methionine (AHA) or azide-modified mannose (ManNaz). 2) The modified amino acids or monosaccharides are incorporated into the newly synthesized matrix. 3) After culture and fixation, a fluorescent dye (DABCO 555) is “clicked” onto the azide groups of the newly synthesized proteins (4) and proteoglycans (5) in a copper-free click chemistry reaction. B) Fluorescently labeled proteins and proteoglycans were imaged with confocal microscopy after 1 day or 7 days of culture. Scale bar = 10 µm. C) Quantification of the area percentage of nascent proteins (left) and nascent proteoglycans (right). Data are represented as mean ± standard deviation. Statistical significance was determined using two-way ANOVA with a Tukey’s multiple comparisons test (∗p < .05, ∗∗p < .01, ∗∗∗p < .001, and ∗∗∗∗p < .0001) (n = 3 replicates).

Specific variations in matrix production by encapsulated hMSCs across different hydrogel formulations and fabrication techniques were observed and quantified by total area percentage, as deposition exceeded the pericellular location after 7 days (**Figure 6C**). First, considering biochemical cues, differences between HA, L/H-dECM, and L/H-Col I hydrogels were evident. After 7 days, L/H-dECM constructs (FLight + bulk) exhibited the highest new ECM secretion, with L-dECM showing ≈20% proteins and ≈25% proteoglycans, and H-dECM with ≈23% proteins and ≈17% proteoglycans. Conversely, L/H-Col I constructs produced the least ECM, with L-Col I showing ≈15% and ≈10% of proteins and proteoglycans, respectively, while H-Col I had ≈14% proteins and ≈16% proteoglycans. HA hydrogels produced a significant number of proteins (≈21%) but a low number of proteoglycans (≈10%). L/H-dECM stimulated protein and proteoglycan production, likely due to the diverse biochemical signals present in dECM hydrogels, as shown in **Figure 2**, compared to the poorer environment in HA and L/H-Col I constructs. [86]

Second, hydrogel architecture also affected total ECM production. FLight hydrogels exhibited more proteins than bulk, whereas bulk constructs possessed an equal number of proteoglycans, on average, compared to FLight. This observation may be explained by the presence of fibrillar collagens, which are restricted in denser hydrogel matrixes (i.e., bulk) compared to proteoglycans. This suggests that proteins and proteoglycans accumulate with different spatial distributions after being secreted by hMSCs within hydrogels, which may be attributed to variations in the overall size and assembly of these molecules, as numerous proteins form large fibrillar structures in contrast to proteoglycans. This aligns with earlier findings that proteoglycans exhibit higher diffusion rates compared to proteins like collagen.[87–89] Note that, in FLight constructs, newly secreted proteins but not proteoglycans were deposited in an aligned fashion that mimicked the cartilage deep zone, likely due to microchannels guiding ECM deposition.

### 2.6. Long-term cartilage tissue maturation

To assess whether the differences in cell shape and early anabolic superiority in the L/H-dECM FLight hydrogels could be maintained, the deposition of cartilage at later time points was characterized. Histological and immunohistological staining was conducted after 56 days of culture (**Figure 7A**). In line with previous literature, Col I content was notably reduced in bulk and FLight samples containing L/H-dECM compared to those with HA alone or in L/H-Col I. Conversely, after 56 days of culture, Col II expression was significantly increased in L/H-dECM constructs for both bulk and FLight samples. Furthermore, GAG staining closely resembled the bovine articular cartilage control (**Figure S6**) when L/H-dECM was employed and was notably enhanced for FLight hydrogels.

**Figure 7.**
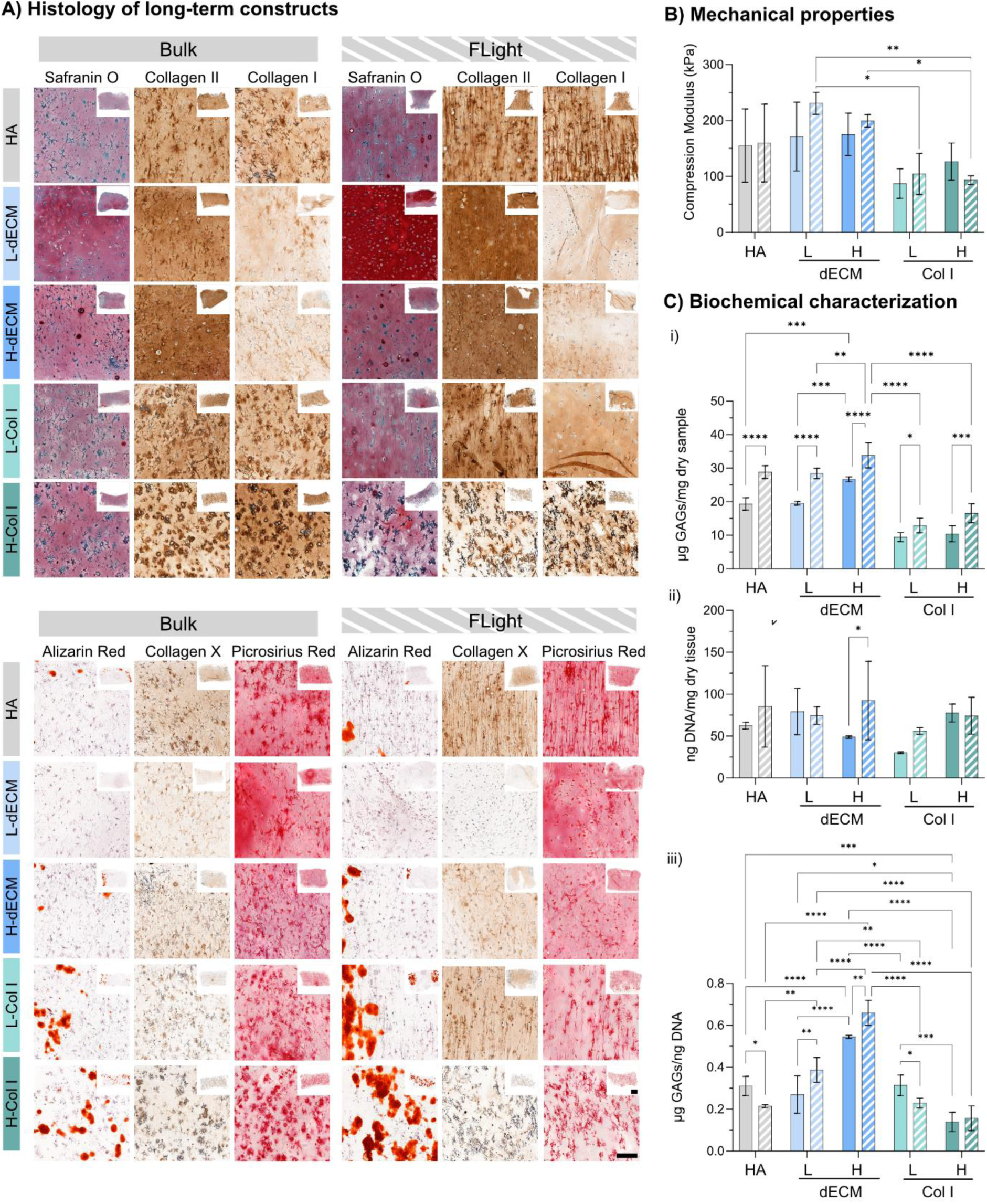
Characterization of neocartilage deposition. A) Histological and immunohistological staining of bulk and FLight gels profile sections over 56 days of culture for GAGs (SafraninO), collagen II, and collagen I. Scale bar of full sample = 1 mm. Scale bar close-up = 100 µm. B) Mechanical properties of the constructs after 56 days. C) GAG quantification (i), DNA quantification (ii) and GAGs/DNA quantification (iii) of hydrogels after 56 days of culture. Data are represented as mean ± standard deviation. Statistical significance was determined using two-way ANOVA with a Tukey’s multiple comparisons test (∗p < .05, ∗∗p < .01, ∗∗∗p < .001, and ∗∗∗∗p < .0001). (n = 3 replicates).

To determine how closely the mechanical properties of the engineered cartilage matched those of native cartilage, samples were subjected to unconfined compression testing. Consistent with the cartilage-like ECM deposition observed in histological staining, the compressive modulus increased throughout the culture period (**Figure 7B**). Starting from relatively soft hydrogels (∼ 1-2 kPa) at day 1, the compression modulus reached values of ∼200 KPa for FLight hydrogels after 56 days for L/H-dECM-composed hydrogels. To our knowledge, achieving high biomechanical properties using hMSCs is rare, often necessitating high initial cell concentrations or the presence of supporting structures.[90]

The levels of GAG content, a crucial component of articular cartilage and an important factor in tissue mechanics, along with cell proliferation, were evaluated after 56 days of culture (**Figure 7C**). Over this period, no significant differences were observed in DNA when normalized to the construct weight between bulk or FLight. However, weight-normalized GAGs showed significant differences at 56 days, always being higher for FLight than bulk hydrogels. When GAG was normalized by DNA content, the significance was maintained for L/H-dECM-containing hydrogels. The GAGs/DNA ratio in the weight-bearing regions of human articular cartilage is approximately 0.2 µg ng^-1^.[91] Our engineered cartilage grafts surpassed this value, indicating that cells were very metabolically active during this period.

Chondrogenic hypertrophy of adult hMSCs, marking the terminal stage of chondrocyte differentiation, is undesirable in cartilage regeneration approaches. This stage is associated with subsequent apoptosis and tissue mineralization.[92,93] hMSCs acquire hypertrophic traits, including upregulation of type-X collagen (COL-X) and deposition of calcified ECM.[94] Additionally, they begin producing ECM-degrading factors such as matrix metalloproteinase 13 (MMP13).[95] Therefore, we further evaluated the expression of such markers (**Figure 7A bottom**). Alizarin red was employed to visualize calcification; after 28 days of culture, no signs of calcification were observed (data not shown). However, after 56 days of culture, calcification was present and more prominent in L/H-Col I. Col X expression after 56 days of culture was less pronounced in L/H-dECM samples compared to others. Finally, samples were stained with Picrosirius red to visualize the total collagen content, as indicated by the red intensity. Both bulk and FLight samples exhibited increased Picrosirius red staining intensity over time, indicating continuous neocartilage deposition by the embedded cells. However, neither of the conditions reached the control intensity, suggesting that further maturation may be necessary to achieve native-like values. In conclusion, both initially rounded (bulk) and elongated (FLight) cells produced more Col II than Col I in L-dECM, indicating that the 3D cell shape did not control the type of collagen produced. Furthermore, the high anabolic response in L-dECM visualized in nascent ECM deposition was maintained over time, as seen by enhanced mechanical properties and GAG production. Chondrogenic hypertrophy was slightly lower in L-dECM bulk and L-dECM FLight hydrogels, highlighting their potential as a promising platform for long-term cartilage tissue maturation.

### 2.7 Short- and long-term deposition of aligned collagen

Finally, the internal collagen structure must resemble articular cartilage architecture. Collagen fibrils provide tensile strength and resistance to deformation in articular cartilage. Thanks to their aligned structure, they contribute to articular cartilage’s overall structural integrity and mechanical properties, allowing it to withstand compressive forces experienced during joint movement.

In this study, we investigated how cells respond to anisotropic matrices compared to bulk over different time frames and secrete aligned proteins. After just 1 day of encapsulation, hMSCs exhibited cell alignment in FLight matrices (**Figure 5**), particularly in those composed of HA and L-dECM, which later facilitated optimal protein deposition (higher percentage of area occupied by newly synthesized proteins, after 7 days of culture, **Figure 6**). Analysis using the orientationJ plugin (ImageJ) revealed distinct protein deposition patterns between bulk and FLight hydrogels after 7 days (**Figure 8A**). Specifically, FLight gels exhibited an even color distribution (blue), indicative of aligned protein deposition, whereas bulk gels showed a wide range of colors, suggesting a lack of alignment. Additionally, FLight gels composed of HA, L-dECM and L-Col I demonstrated a single peak in the orientation distribution graphs, while all other conditions showed an even distribution at different angles. FLight gels for H-dECM and H-Col I showed no alignment, suggesting that they may hinder the presence of microchannels and hamper the aligned deposition of proteins. Furthermore, cells in bulk gels did not have a preferred direction for protein deposition. Notably, protein deposition in FLight hydrogels occurred on both edges of cell polarization, indicating that protein alignment is independent of cell migration but rather dependent on void space localization.

**Figure 8.**
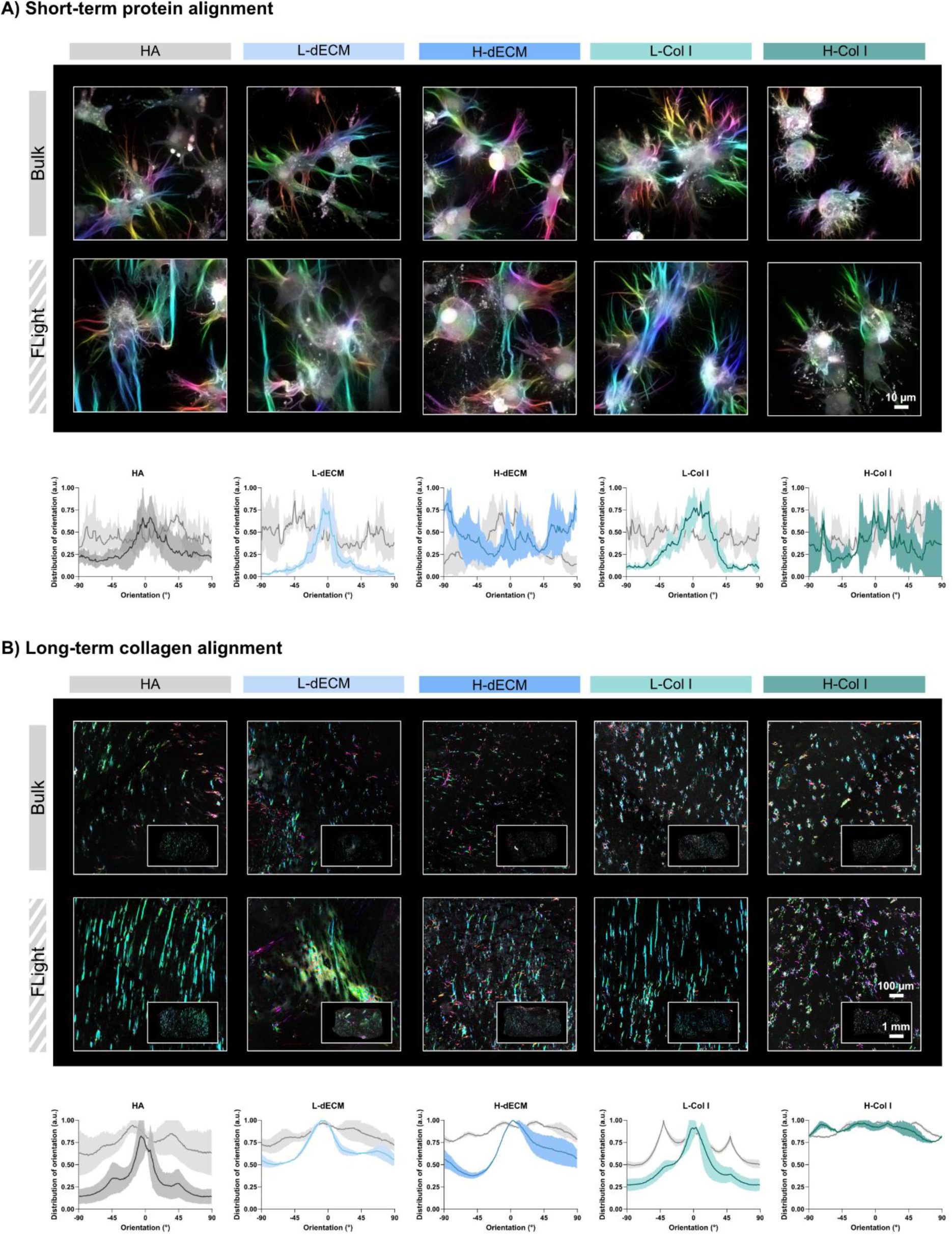
Evaluation of collagen alignment after short- and long-term culture. A) Nascent protein labeling after 7 days in culture, color-coded by angle. Scale bar = 10 µm. Below is the orientation distribution plot of FLight (colored) vs Bulk (grey). B) Angle color-coded images from the polarized microscope after staining with picrosirius red. Scale bar for full sample = 1 mm. Scale bar for close-up = 100 µm. Below is the orientation distribution plot of FLight (colored) vs Bulk (grey). Data are shown as means ± SD; three independent experiments; n = 3.

To visualize whether this structure was maintained over long culture periods, Picrosirius red slides were observed under the polarized-light microscope (PLM) (**Figure 8B**). While bulk samples lacked distinct collagen orientation, a clear alignment was evident in the anisotropic FLight hydrogels, particularly noticeable after applying the orientation color map (**Figure 8B**). This color map enabled quantification of collagen fibril coherency and orientation (**Figure 8B**), revealing a significant difference between the engineered constructs. Specifically, the fibril coherency values for HA, L-dECM, H-dECM, L-Col I, and H-Col I were 0.2±0.053, 0.16±0.005, 0.047±0.008, 0.139±0.034, and 0.112±0.013 for bulk hydrogels, and 0.568±0.086, 0.285±0.048, 0.212±0.046, 0.603±0.069, and 0.107±0.016 for FLight hydrogels, respectively. FLight demonstrated significantly higher fibril coherency, with predominant alignment observed in HA, L-dECM, and L-Col I. Despite increased light scattering associated with dECM incorporation, fiber alignment resembling the articular cartilage deep zone was achieved.

## Conclusion

In this study, we successfully employed FLight biofabrication to produce cell-laden anisotropic hydrogels using biologically relevant resins previously unexplored with this method, such as articular cartilage dECM and Col I. This innovative technique enabled the precise printing of challenging resins with high resolution while simultaneously mitigating the immune response. Furthermore, our findings highlight the importance of studying early cell-biomaterial interactions, including changes in cell morphology and nascent ECM deposition, which significantly influence long-term tissue maturation. Notably, we demonstrated that an initial elongated morphology does not dictate the type of matrix produced. Specifically, we found that regardless of initial cell shape, when encapsulated in articular cartilage dECM-containing matrices and cultured in chondrogenic media, hMSCs deposited more Col II than Col I and produced high amounts of GAGs. Furthermore, we found that the presence of dECM contributes mildly to the protection against hypertrophy when compared to HA or Col I. In summary, our findings suggest that FLight-fabricated hydrogels can better orient the produced ECM compared to bulk and that the use of L-dECM allows better cartilage maturation in terms of mechanical properties, GAG production and Col II deposition. Therefore, anisotropic hydrogels based on L-dECM significantly enhance articular cartilage tissue maturation, closely approaching native composition, structure, and biophysical properties.

### Experimental Section

HA was purchased from HTL Biotechnology. Purified Type I Collagen of bovine origin was obtained from Symatese. All other chemicals were purchased from Merck, and cell culture reagents were purchased from Gibco unless indicated otherwise.

#### Synthesis of hyaluronic acid tyramine

HA tyramine was synthesized using a previously reported protocol[96] with some modifications. Sodium hyaluronate (1.0 g, 2.5 mmol of COOH groups) was dissolved in 200 mL of MilliQ-water overnight with gentle stirring. EDC (0.95 g, 5 mmol) and NHS (1.15 g, 10 mmol) were added, and the solution was stirred for 30 min at room temperature. The aqueous solution of Tyr-HCL (0.86 g, 5 mmol) was added to the mixture to initiate the reaction; the pH value was controlled at 4.75, and the reaction was carried out overnight. Then the pH value was brought to 7.0 by adding 1 M NaOH solution to terminate the reaction. The product was purified by dialysis for 7 days against distilled water to remove residual unreacted reagents. The resultant HA tyramine conjugate was collected as white foam after freeze-drying. The degree of substitution (DS), defined as the number of substituted groups per 100 disaccharide units, was calculated from ^1^H-NMR (D2O) spectral data (**Figure S1**,) by comparing the integral values of the aromatic protons of tyramine (peaks at 6.86 and 7.17 ppm) and the methyl protons of HA (1.9 ppm). HA tyramine with a degree of substitution (DS) of 8.8% was used in this study.

#### Preparation of articular cartilage dECM

Bovine knees were acquired from a local butcher (Angst AG Zurich). The knees were opened to access the articular cartilage, which was then peeled off using a blade. The cartilage pieces were collected into falcon tubes filled with a 5% phosphate buffer saline (PBS)-antibiotics mixture (penicillin-streptomycin, 15070063, Gibco). The larger pieces of cartilage were washed three times with PBS, minced into small pieces using a blade, and then put back into PBS to avoid drying. To achieve physical decellularization, the cartilage pieces were frozen with liquid nitrogen and then thawed at 37 °C, which corresponds to one cycle. Three freeze-thaw cycles were conducted in total. The articular cartilage was then put in a 0.1 M HCl solution (10 mL g^−1^ of wet cartilage) for 24 h with 125 rpm stirring at 30 °C using a water bath. A low concentration of HCl is used to prevent damage to collagen fibers, as it can disrupt ionic intramolecular bonds, leading to ECM deterioration. Subsequently, the cartilage was put in a sieve and washed three times with deionized water. After the washing steps, the dECM was placed into falcon tubes, frozen with liquid nitrogen, and lyophilized to dry the decellularized cartilage. The dried material was cryomilled (CryoMill, Retsch GmbH, verder scientific) with two cycles of 2.5 min duration each at 25 Hz, resulting in a powder of dECM. Then, a pepsin solution was prepared by fully dissolving 1 mg mL^−1^ pepsin from porcine gastric mucosa (lyophilized powder >2500 units mg^−1^, Sigma) in 0.1 M HCl solution, as previously described.[97] The dECM powder was then put into the pepsin solution (10 mgmL^−1^) and left under stirring at 200 rpm at RT for 48 h. Afterwards, neutralization of the dECM was conducted. At 1h before starting neutralization, the digestion vial and the solutions needed (PBS, NaOH, and HCl) were placed on ice. Then, the stir plate was moved up to 350 rpm and 1/20 of the starting volume was added of 1M NaOH. There was a waiting time of 10 min before measuring the pH. Depending on the pH, either 1M NaOH or 1M HCl were added to titrate the pH to 7.4. Later, 1/9 of the total liquid volume of 10x PBS was added. Finally, the neutralized solution was lyophilized. The dry material was stored in the -80° freezer.

#### Sample preparation for biochemical assays

To prepare the dECM and cartilage samples for characterization, papain solution was prepared, containing 10 mM L-cysteine HCl (C-7477, Sigma), 100 mM sodium phosphate (567547, Merck), and 10 mM Ethylenediaminetetraacetic acid (EDTA, A2937, AppliChem) in MilliQ-water. To achieve a pH of 6.3, the solution was adjusted with 1 M HCl. 270 μg mL^−1^ of papain was added right before using the digestion solution. Then, 500 μL were added to approximately 10 mg of sample material (the exact quantities were noted, and the concentrations were calculated accordingly). The solutions were digested at 60 °C overnight with 1000 rpm shaking.

#### DNA quantification

The amount of DNA was quantified using a Quant-iT PicoGreen dsDNA assay kit (Invitrogen) following manufacturer’s protocol. Briefly, 1X Tris-EDTA (TE) Buffer was prepared by diluting 1:20 the 20X TE buffer from the kit in MilliQ-water. The PicoGreen was then diluted 1:200 in the 1X TE buffer. DNA standard solutions with a final concentration of 1000, 500, 100, 10, and 0 ng mL^−1^ were prepared diluting the 100 μg mL^−1^ DNA standard from the kit into 1x TE buffer. Then, 10 μL of the digested samples of cartilage and dECM of the papain solution as background were distributed into a 96-well plate. These were diluted with 40 μL 1X TE buffer each. The DNA standards were placed on the plate with 50 μL each. In all samples 50 μL of PicoGreen solution were added, reaching a total dilution of 1:10. Duplicates for all samples were conducted. The fluorescence was measured with a microplate reader (Synergy H1 Hybrid Reader, BioTek) at 480 nm excitation and 520 nm emission. The resultant data was analyzed with Excel to obtain a standard curve, from which the concentration of DNA content in the tested samples was calculated.

#### GAG quantification

GAG content was quantified using Blyscan sGAG assay according to the manufacturer’s instruction. First, GAG standards were prepared containing 0.0, 1.0, 2.0, 3.0, 4.0 and 5.0 µg of the reference standard and mixed with the digestion buffer (without papain). Test samples of cartilage were diluted 1:200, whereas the dECM was diluted 1:100. Then, the Blyscan dye reagent was added to each tube and the tubes were placed on a mechanical shaker for 30 min at 100 rpm. During this time, a sulphated GAG-dye complex formed and precipitated from the soluble unbound dye. Later, the tubes were transferred to a centrifuge and spun down at 12000 rpm for 10 min. The tubes were then inverted and drained by gently tapping them on a tissue. The dissociation reagent was added in a volume of 0.25 mL to each tube and vortexed to release the bound dye into solution. 100 μL of each sample were transferred to individual wells in a 96-well plate by non-rapid pipetting, to avoid foam formation, which could cause abnormal absorbance readings. The measurement was performed in a microplate reader (Synergy H1 Hybrid Reader, BioTek) with 656 nm absorbance. The resultant data was analyzed with Excel to obtain a standard curve, from which the concentration of GAG content in the tested samples was calculated.

#### Sample preparation for proteomic analysis

Two milligrams of lyophilized material from each sample was homogenized in 200 μL mg^-1^ of 6 M guanidine hydrochloride (Gnd-HCl), 100 mM ammonium bicarbonate (ABC) at power 8 for 1 minute (Bullet Blender, Model BBX24, Next Advance, Inc.) and vortexed (power 5) at room temperature overnight. Homogenate was spun at 18,000 x g (4 °C) for 15 min and the supernatant was collected as the soluble ECM (sECM) fraction. Pellets were then treated with freshly prepared hydroxylamine buffer (1 M NH2OH−HCl, 4.5 M Gnd−HCl, 0.2 M K2CO3, pH adjusted to 9.0 with NaOH) at 200 μL mg^-1^of the starting tissue dry weight. Samples were homogenized at power 8 for 1 minute and incubated at 45 °C with shaking (1000 rpm) for 4 h. Following incubation, the samples were spun for 15 min at 18,000 x g, and the supernatant was removed and stored as the insoluble ECM (iECM) fraction at −80°C until further proteolytic digestion with trypsin. All fractions were subsequently subjected to enzymatic digestion with trypsin using a filter-aided sample preparation (FASP) approach and desalted during Evotip loading (described below).

#### LC-MS/MS analysis

Digested peptides were loaded onto individual Evotips following the manufacturer’s protocol and separated on an Evosep One chromatography system (Evosep, Odense, Denmark) using a Pepsep column (150 µm inter diameter, 15 cm) packed with ReproSil C18 1.9 µm, 120Å resin. Samples were analyzed using the instrument’s default “30 samples per day” LC gradient. The system was coupled to the timsTOF Pro mass spectrometer (Bruker Daltonics, Bremen, Germany) via the nano-electrospray ion source (Captive Spray, Bruker Daltonics). The mass spectrometer was operated in PASEF mode. The ramp time was set to 100 ms and 10 PASEF MS/MS scans per topN acquisition cycle were acquired. MS and MS/MS spectra were recorded from m/z 100 to 1700. The ion mobility was scanned from 0.7 to 1.50 Vs/cm2. Precursors for data-dependent acquisition were isolated within ± 1 Th and fragmented with an ion mobility-dependent collision energy, which was linearly increased from 20 to 59 eV in positive mode. Low-abundance precursor ions with an intensity above a threshold of 500 counts but below a target value of 20000 counts were repeatedly scheduled and otherwise dynamically excluded for 0.4 min.

#### Global proteomic data analysis

Data was searched using MSFragger via FragPipe v21.1. Precursor tolerance was set to ±15 ppm and fragment tolerance was set to ±0.08 Da. Data was searched against UniProt restricted to *Bos taurus* with added common contaminant sequences (37,606 total sequences). Enzyme cleavage was set to semi-specific trypsin for all samples. Fixed modifications were set as carbamidomethyl (C). Variable modifications were set as oxidation (M), oxidation (P) (hydroxyproline), dioxidation (P), deamidation (NQ), Gln->pyro-Glu (N-term Q), and acetyl (Peptide N-term). Label-free quantification was performed using IonQuant v1.10.12 with match-between-runs enabled and default parameters. Soluble and insoluble ECM fractions were searched separately and merged after database searching. Results were filtered to 1% FDR at the peptide and protein level.

#### Cell isolation and expansion

Human bone marrow was obtained as surgical waste material with the written consent of the patients. The Swiss Human Research Act does not apply to research that involves anonymized biological material and/or anonymously collected or anonymized health-related data. Therefore, this project does not need to be approved by the ethics committee. General consent, which also covers anonymization of health-related data and biological material, was obtained. MSCs were isolated via density centrifugation separation and plastic adhesion. Isolated cells were plated at a concentration of 3000 cells cm^−2^ and expanded in minimal essential medium (MEMα) (without nucleosides, Gibco), supplemented with fetal bovine serum (FBS, 10%, Gibco), gentamicin (100 µgmL^−1^, Gibco), and fibroblast growth factor 2 (FGF-2, 5 ng mL^−1^, PeproTech). Cells were cultured at 37 °C in a humidified atmosphere at 5% CO2. Cells were passaged until they reached 80% confluency. At passage 3, cells were trypsinized, combined with DMEM, 10% v/v FBS, and 10 µg mL^−1^ gentamicin and collected by centrifugation (5 min, 500 rcf). Finally, the cell pellet was carefully mixed with the different hydrogel precursors at a concentration of 5 million cells mL^-1^. After crosslinking, constructs were kept in chondrogenic media; Glutamax DMEM supplemented with 10 ng mL^−1^ transforming growth factor β3 (TGF-β3, Peprotech), 50 μg mL^−1^ L-ascorbate-2-phosphate, 40 μg mL^−1^ L-proline, 50 μg mL^−1^ gentamicin (Gibco), and 1% ITS+ Premix (Corning).

#### Preparation of Photoresins

At first, stock solutions were prepared in the following way. HA, dECM, and Col I were dissolved for all experiments in PBS. HA tyramine was dissolved to achieve a stock concentration of 1.5% w/v and left at RT for over 24 h on a shaking plate covered with aluminum foil. Stock solutions of 20 mg mL^-1^ of dECM and Col I were prepared by dissolving them in PBS and putting them in the fridge for 24h. The stock solutions of the photoinitiators were freshly prepared every time to obtain 10 mM and 100 mM of Ru/SPS (Advanced Biomatrix), respectively. They were kept closed with aluminum foil to avoid light exposure. The photoresins were then prepared by mixing the required volume of stock solutions (2.5 dECM: 0.8% HA 2.5mg mL^-1^ dECM, 5 dECM: 0.8% HA-TYR 5mg mL^-1^ dECM, 2.5 Col I: 0.8% HA 2.5mg mL^-1^ Col I, 5 Col I: 0.8% HA-TYR 5mg mL^-1^ Col I) with PBS. After each component was added, vortexing and pipetting up and down were important to ensure a good mixture. Also, centrifugation for a few seconds was carried out to remove air bubbles. As the stock solutions and final photoresins are viscous, positive displacement pipettes are used to handle them.

#### FLight biofabrication

The (bio)photoresins were pipetted into PDMS rings (2 mm height, 4 mm diameter) or 1.5 ml Eppendorf cup tube (2.5 mm height, 10 mm diameter), generating hydrogels with a volume of 25 μL or 200 μL, respectively, which were later placed into wells of a 24- or 12-well plate, respectively. FLight projections were performed in a custom-made FLight printer deploying a fiber coupled laser (405 nm, 1000 mW power) with a high speckle contrast ratio (to achieve the modulation instability necessary for microfilament formation). The light was expanded (using a 4f magnification system) to cover a digital micromirror array device (DMD; DLP6500FYE, Texas instruments) with a pixel resolution 1920×1080 and a pixel pitch 7.56-µm. The DMD was used to shape the light to the desired cross-section image, which was then expanded using a set of plano convex lenses (1:1 magnification) with an aperture in between (to isolate the reflected image from the diffraction pattern of the DMD), and the image projected in a top-down orientation. The exposure time was 8s, and a light intensity of 62.5 mW cm^-2^ was used. After printing, uncrosslinked (bio)photoresin was removed with PBS prewarmed to 37 °C. The washing was repeated three times until the hydrogels became transparent, indicating that the Ru/SPS had been removed.

#### Bulk biofabrication

The bulk samples were crosslinked for 8s in a UV box with a tunable light intensity (35% dimming), which was adjusted to match the light dose of the FLight procedure. This method will be referred to as the bulk method.

#### Assessment of Microstructure

Printed acellular hydrogels were fabricated as previously described using the FLight or LED method in PDMS rings with the addition of 1 μg mL^-1^ acryloxyethyl thiocarbamoyl rhodamine B (Rhodamine B acrylate) (25404-100, PolySciences) in dimethyl sulfoxide (DMSO). The hydrogels were then kept in PBS to avoid dehydration. Imaging was performed with a confocal microscope (Leica TCS SP8, Leica). For the constructs, a z-stack of 100 μm (0.5 z-step) was performed to obtain the diameter of the filaments, which were manually measured with ImageJ.

#### Rheological properties

Gel precursors were tested on a rheometer (Modular Compact Rheometer MCR302e, Anton Paar) using a PP10 plate and a wavy glass plate on the bottom to avoid slippage. The shear-strain (oscillating) measurement was performed over 20 min at 1 Hz frequency and 1% strain with a gap distance of 0.5 mm and a temperature of 25 °C. To measure the baseline, during the first 5 min the resins were not exposed to any light. After 5 min the light source was turned on and for 15 min the resin was exposed to the light at 60% intensity. Storage and loss modulus (G’ and G”) were measured over the whole 20 min, when the storage modulus approaches a plateau. Each sample was loaded with 45 μL gel solution and put in the center of the glass plate surrounded by silicon oil (Silitech SA) and a wet tissue to prevent dehydration. All measurements were performed with freshly prepared resins. For each tested gel composition, three measurements were taken. The obtained G’ and G” as well as the stiffness were plotted and compared across the different resin compositions. Shear stress relaxation measurements were conducted consecutively to the rheological tests without a time interval between the two tests. The crosslinked gels were incubated at 37 °C for 30 min without light with 1 Hz frequency and 1% strain to allow fibrils formation. Afterwards, the hydrogels were constantly compressed to 10% strain under stress relaxation mode for 45 min and the stress was recorded as a function of time. The stress relaxation time τ½ was calculated as the time in which the initial stress of the hydrogel relaxed to ½ of its initial value. Three measurements were taken for each gel composition tested.

#### Uniaxial compression tests

A compression test of printed acellular constructs crosslinked via Flight or LED method was performed with a texture analyzer machine (TA.XT.plus Texture Analyser, Stable Micro Systems). A 5 N load cell was used with a test speed at 0.01 mm s^-1^ and 15% strain. All tests were performed on wet hydrogels at RT. Before testing, a picture of each sample was taken to measure the exact diameter with ImageJ and then be able to adjust the obtained results. For each tested gel composition, three measurements were taken. The compressive moduli were calculated as the slope from 0% to 3% strain and compared across the different gel compositions.

#### Swelling properties

To determine the swelling ratio of the hydrogels, the photoresins were crosslinked by the FLight or LED method (V=200 μL). The gels were then carefully washed 3x in PBS to remove the uncrosslinked resin. Afterwards, the gels were collected to measure the first time point weight following crosslinking, and the excess water was removed with tissue paper (Kimteck). The scaffolds were then incubated in 4 mL of PBS in a 12-well plate at 37°C and weighed after 24h of fabrication, removing again the excess water with tissue paper. The swelling ratio was determined as the ratio of the hydrogel mass after 24 h divided by its initial mass.

#### Hydrogel degradation

To assess the degradation properties of the scaffolds, cast hydrogels were incubated at 37 °C in an enzyme solution consisting of 1 U mL^-1^ hyaluronidase (type I-S from bovine testes, Sigma Aldrich) and 0.3 U mL^-1^ collagenase (type II from Clostridium histolyticum, Sigma Aldrich) in PBS. These concentrations were adapted from O’Shea and colleagues[98]. The scaffolds were taken out of the enzyme solution and weighed at regular intervals. The degradation ratio was calculated by dividing the mass of the hydrogel at a given time point by the mass of the hydrogel at the start of the experiment (after 24 h of swelling).

#### Printability

The projection image of a snowflake was created using Affinity Designer 2.3 with a fixed resolution of 1920×1080 pixels, with each pixel equal to about 7.3 μm. The white colored image corresponded to 100% light intensity (≈62.5 mW cm^-2^). Photoresins were prepared as previously described, adding 1μg mL^-1^ rhodamine B acrylate in DMSO. The image was then exported as PNG file and projected with the custom-made FLight printer in rectangular wells of a glass cuvette containing 250 μL of resin each. The well was then washed 3 times with PBS to remove the uncrosslinked photoresin. Imaging was performed with a confocal microscope (Leica TCS SP8, Leica), and ImageJ was utilized for the posterior analysis.

#### Culture of immune cells

Cells were cultured in suspension in T-150 cm^2^ cell culture flask at a subculture density of 1–5 × 10^5^ cells mL^-1^. Cells were cultured in RPMI-1640 medium supplemented with 10% FBS, 1% penicillin-streptomycin, 1% GlutaMAX 100-X (Gibco), 1% MEM Non-Essential Amino Acids Solution 100-X and 1% sodium pyruvate at 37 °C, 5% CO2 and 95% humidity. To differentiate THP1 cells into adherent and nonpolarized M0 macrophages, cells were seeded as 1×10^5^ cells cm^-2^ in well plates and incubated with RPMI-1640 medium supplemented with 5 ng mL^-1^ phorbol myristate acetate (PMA) for 24 h. Afterward, the medium was aspirated, and the adherent macrophages were washed twice with PBS and cultured for 48 h in a PMA-free medium. Cells were cultured on 65 μl hydrogels at a density of 6.5×10^4^ cells mL^-1^.

#### Live/Dead and SHG Imaging

Samples were biofabricated as previously described and incubated in DMEM supplemented with 1:1000 CalceinAM (Invitrogen) and 1:500 Propidium Iodide (PI, Fluka) for 40 min without FBS. Imaging was then performed on a Leica SP8 microscope (Leica) equipped with a 25× objective. SHG imaging was collected at ≈440 nm with 880 nm two-photon excitation (Mai Tai laser, Spectra-Physics). Z-stacks were acquired from the sample surface at 2 µm steps and 100 µm into the sample. The pictures reported in the work resulted from maximum intensity z-projection.

#### Nascent Matrix Labeling and Quantification

The nascent ECM labeling protocol was adapted from Loebel and colleagues.[99,100] hMSCs were encapsulated at a density of 5×10^6^ cells mL^-1^ in hydrogels (25 μl) and cultured in 48-well plates at 37 °C, 5% CO2 and 95% humidity. The cell-laden hydrogels were cultured in chondrogenic media (DMEM 21013024, 1% ITS+, 10 μg mL^-1^ gentamycin, 50 μg mL^-1^ ascorbic acid, 40 μg mL^-1^ L-proline, 40 μg mL^-1^ dexamethasone, and 10 ng mL^-1^ TGF-*β*3). The chondrogenic media was further supplemented with 2 mM glutamine, 1 mM sodium pyruvate, and 0.201 mM L-cysteine. To obtain the L-Azidohomoalanine (AHA) media, 100 μM AHA and 101 μM L-methionine were added. To obtain the N-azidoacetylmannosamine-tetraacylated (ManNAz) media, 50 μM NMAM and 202 μM L-Methionine were added instead. The media was exchanged every 3 days. At different time points (day 1 and day 7) of maturation, hydrogels were first washed three times in 1% bovine serum albumin (BSA) in PBS, followed by a 45-min incubation at 37 °C in a fluorophore-conjugated cyclooctyne (DBCO-555) containing solution of 1% BSA in PBS. Then, after three washes in 1% BSA in PBS, hydrogels were fixed in 4% paraformaldehyde (PFA) for 20 min at RT followed by three washes in PBS and storage at 4 °C. The fixed hydrogels were then stained with a plasma membrane stain (CellMask Deep Red, 1:1000 dilution, Invitrogen) and a nuclear stain (Hoechst 33342, 5 μg mL^-1^, Thermo Fisher) for 40 min and were subsequently washed three times with PBS. Imaging was performed with a confocal microscope (Leica TCS SP8, Leica) to acquire z-stacks of 100 μm (0.5 z-step) of nuclei, cell membranes, and labeled matrix, with a 63x magnification. Single cells were cropped from the image to measure the pericellular matrix thickness, and the cell membrane and nucleus area were subtracted. Analysis of the protein’s orientation was carried out using the OrientationJ plugin in ImageJ.[101]

#### Fluorescent confocal microscopy of F-actin cytoskeleton

F-actin cytoskeleton was visualized 1 day after gelation with a confocal microscope (Leica TCS SP8, Leica). Hydrogel constructs were prepared for fluorescent staining by first fixing hMSCs with 4% PFA for 2 hours and permeabilized with 0.5% Triton X-100 in 1× PBS for 5 min. Hydrogel constructs were then stained with Alexa Fluor 565 phalloidin (5µg mL^-1^; Sigma-Aldrich, USA) for F-actin cytoskeleton visualization together with Hoechst 33342 (5 μg mL^-1^, Thermo Fisher) for 40 min for cell nucleus. All samples were rinsed with PBS to remove residual waste and submerged in 1× PBS during imaging. Analysis of cell filopodia was done with FiloQuant plugin.[102] Cell area and cell AR were measured on single cells and analyzed with the particle analysis function of ImageJ. Optical image stacks of single cells were obtained with a 0.5-μm z-axis interval.

#### Histology and Immunohistochemistry

Samples were washed in PBS and fixed in 4% PFA for 2 h. They were then dehydrated in graded ethanol solutions and embedded in paraffin. Sections of 5 μm were cut with a microtome and mounted on glass slides. For the stainings reported below, samples were deparaffinized and rehydrated before staining.

For Safranin O staining, samples were first stained in Weigert’s Iron Hematoxylin solution for 5 min, followed by washing in deionized water and 1% acid-alcohol for 2 s. Sections were washed again in deionized water, stained in 0.02% Fast Green solution for 1 min, and rinsed with 1% acetic acid for 30 s. Finally, sections were stained in 1% Safranin O for 30 min, dehydrated with xylene, and mounted. For alizarin red, samples were stained with 2% alizarin red S in distilled water (pH adjusted to 4.1-4.3 with 10% ammonium hydroxide). Slides were then blotted, rinsed with 95% ethanol, dehydrated to xylene, and mounted. For picrosirius red staining, samples were first stained in Weigert’s Iron Hematoxylin solution for 5 min, followed by washing in deionized water. Slides were then incubated for 1 h in 0.1% of Direct Red 80 in a saturated solution of picric acid. Finally, slides were washed in acidified water, dehydrated to xylene, and mounted.

For immunohistochemistry (collagen I, collagen X, and collagen II staining), antigen retrieval was performed with hyaluronidase (1 % wt/v) at 37 °C for 30 min. Sections were then blocked with 5% bovine serum albumin (BSA) in PBS for 1 h and incubated overnight with the primary antibody, mouse anti-collagen 1 (1:1000, Ab6308, abcam), mouse anti-collagen 2 (1:20, Hybridoma Product II-II6B3, DSHB), or mouse anti-collagen X (1:500) in 1% BSA in PBS at 4 °C. Sections were then incubated with the secondary antibody, goat anti-rabbit IgG-HRP (1:1000, ab6789, abcam), in 1% BSA in PBS for 1 h and developed with the DAB substrate kit (ab64238, abcam) according to manufacturer’s protocol for 5 min. Sections were stained with Weigert’s iron hematoxylin (Thermo Fisher Scientific) for 3 min, destained in 1% acid-alcohol, blued in 0.1% Na2CO3, and finally dehydrated with xylene and mounted.

Slides were then imaged using a 3DHistech slide scanner, while sections stained with picrosirius red were imaged using a Zeiss (AxioImager.Z2) polarised light microscopy. Analysis on the fiber orientation and coherency was carried out using the OrientationJ plugin in ImageJ software.[101]

#### Uniaxial compression tests on cultured samples

Uniaxial unconfined compression tests were performed on a TA.XTplus Texture Analyzer (Stable Micro Systems) equipped with a 500 g load cell. Samples were placed between the compression plates and pre-loaded (2 g on day 56) to ensure full contact with the plates. Samples were allowed to relax for 2 minutes and then were compressed to 15% strain at 0.01 mm s^-1^. The compressive modulus was calculated by linear-fitting the stress-strain curve from 0.5 to 3% strain.

#### Statistical analyses

Analyses were performed in GraphPad Prism 10.2.0. For the comparison of only two groups, unpaired Welch’s t test was used. If more than two groups were compared, depending on the number of variables, ordinary One-way or Two-way analysis of variance (ANOVA) was used, followed by Tukey’s test for multiple comparisons. The fitting of a linear function to the obtained data from the compression test measurements was performed in Excel, and the slope from the equations was used for the statistical analysis. Further, p values are indicated by * (<0.05), **(<0.01), *** (<0.001), and ∗∗∗∗p < .0001). Data is shown as mean ± SD.

## Supporting Information

Supporting Information is available from bioRxiv.

## Acknowledgements

The authors would like to thank the Swiss National Science Foundation for providing financial support (Grant No. SNF IZSEZ0-209669 to MZW). The authors acknowledge the ETH ScopeM imaging facility for their assistance and would like to thank Dr. Philipp Fisch for his help setting up the rheology measurements and Hao Liu for his contributions to FLight technology development.

## Data Availability Statement

The data of this manuscript is available at the ETH Zurich Research Collection: http://hdl.handle.net/20.500.11850/681123.

