## Supporting information for "Anisotropic Articular Cartilage Biofabrication based on Decellularized Extracellular Matrix"

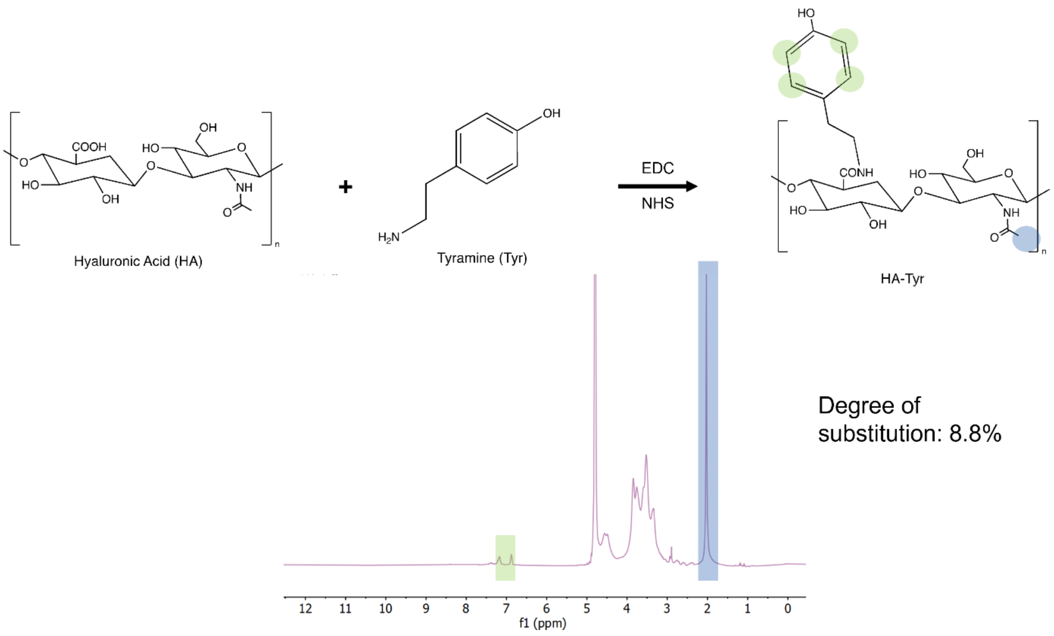


Figure S1. Chemical modification of hyaluronic acid with tyramine. Bottom $^{1}$H NMR spectrum of HA tyramine with a DS of 8.8%.

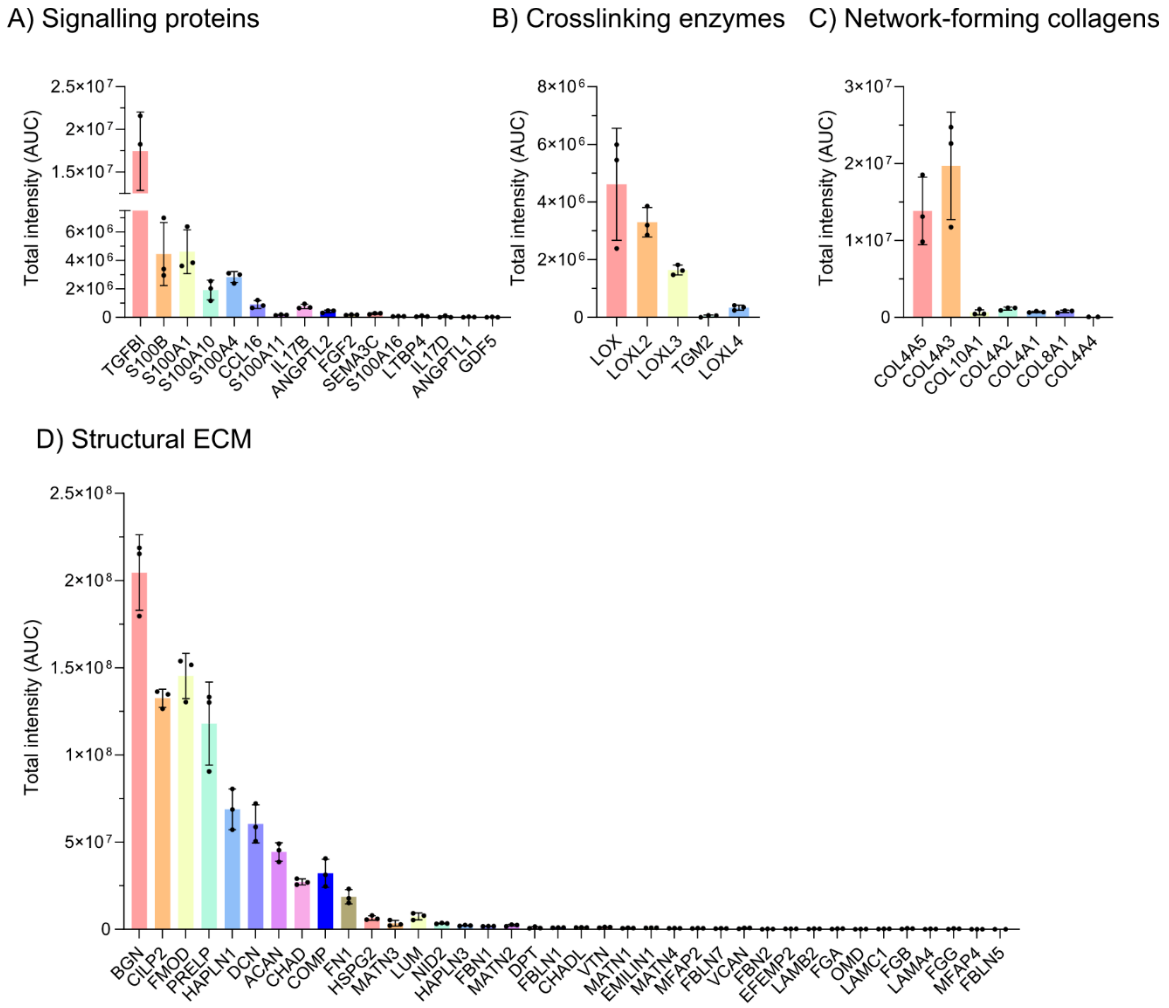


Figure S2. Proteomic analysis of articular cartilage dECM. A) signaling proteins, B) crosslinking enzymes, C) network-forming collagens, and D) structural ECM.

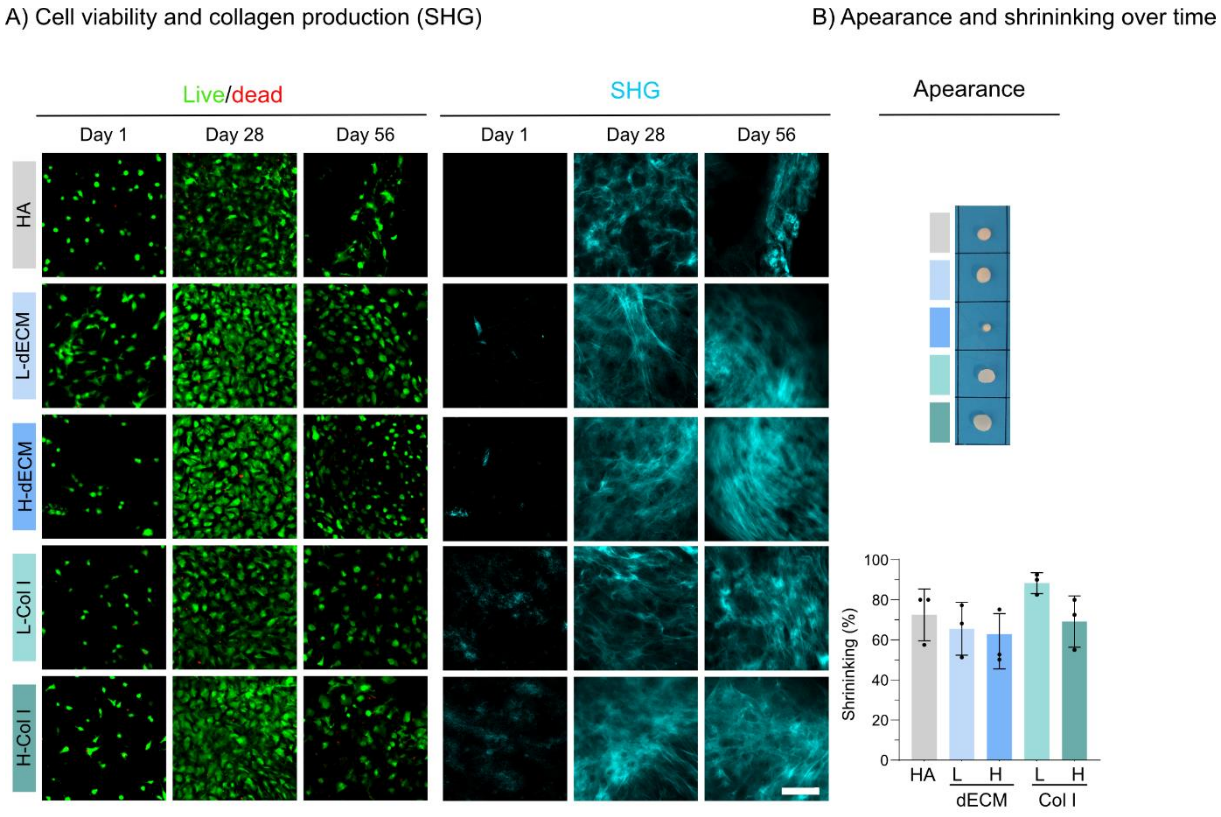


Figure S3. Cell viability (left) and SHG signal (middle) after days 1, 28, and 56 of culture in chondrogenic media for FLight printed gels. Scale bar = 50 µm (A). On the right, there is a picture of the gels after being cultured for 56 days in chondrogenic media and the shrinking measurement (B).

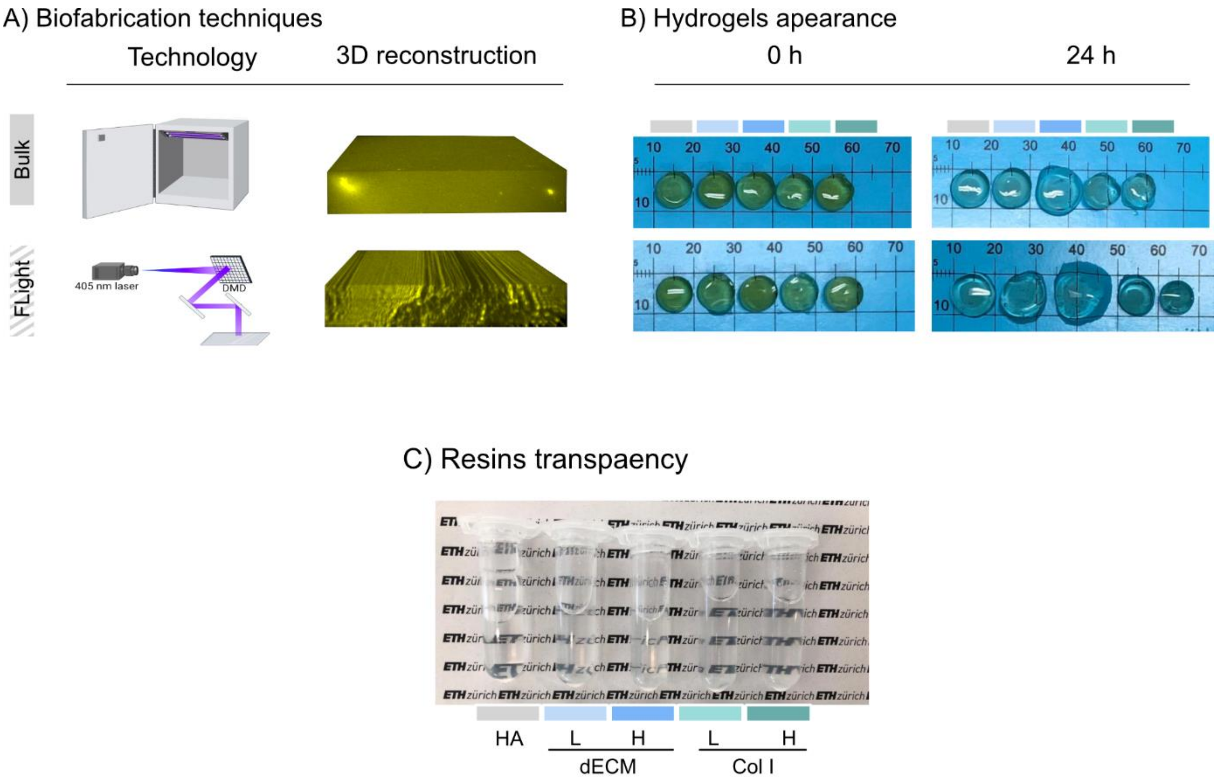


Figure S4. Characterization of the fabricated hydrogels and resin precursors. A) Graphical overview of the two explored technologies with a 3D reconstruction of the printed hydrogels. B) Appearance of the round hydrogels at two different time points: 0h and 24h in PBS, grey= HA, light blue = L-dECM, blue = H-dECM, light green = L-Col I, green = H-Col I. C) Assessment of the resins' transparency.

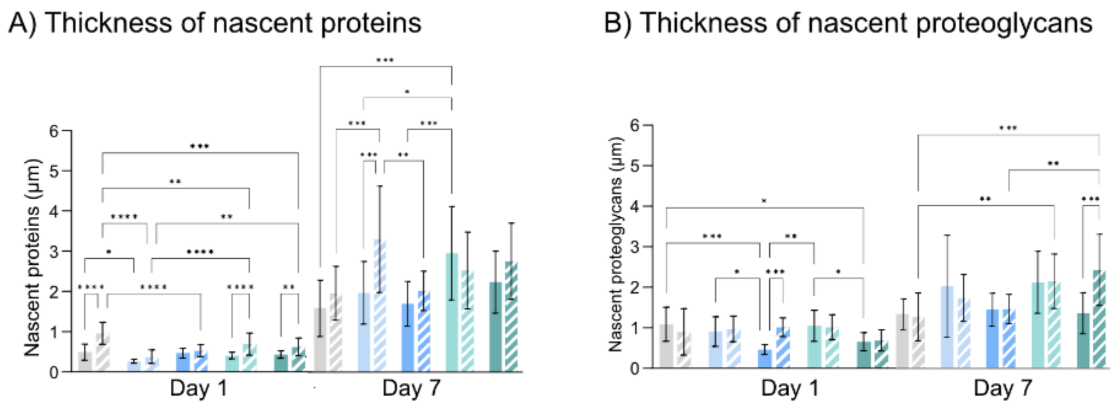


Figure S5. Quantification of the average thickness of nascent proteins (A) and nascent proteoglycans (B), grey= HA, light blue = L-dECM, blue = H-dECM, light green = L-Col I, green = H-Col I, plain filling corresponds to bulk hydrogels and striped to FLight hydrogels. Data are represented as mean ± standard deviation. Statistical significance was determined using two-way ANOVA with a Tukey's multiple comparisons test ($*p < .05$, $**p < .01$, $***p < .001$, and $****p < .0001$). (n = 3 replicates).

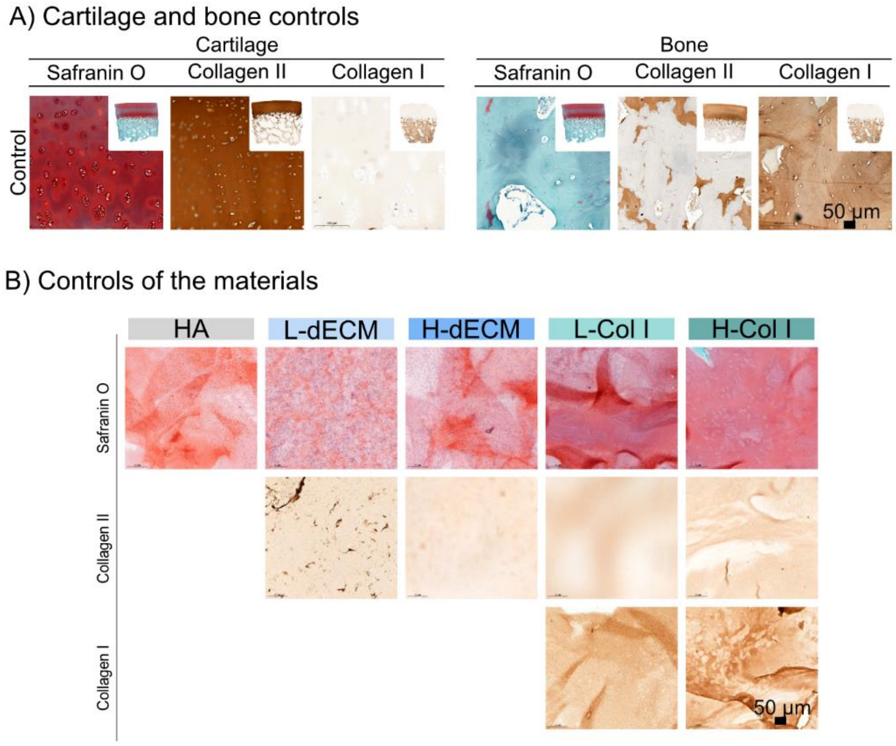


Figure S6. Histology controls. A) Staining for Safranin O, Col II, Col I on bovine cartilage and bone. B) Staining for Safranin O, Col II, Col I on empty gels. Scale bar = 50 µm.
